# Human periventricular nodular heterotopia shows several interictal epileptic patterns, associated with hyperexcitability of neuronal firing

**DOI:** 10.1101/816173

**Authors:** Valerio Frazzini, Stephen Whitmarsh, Katia Lehongre, Pierre Yger, Jean-Didier Lemarechal, Bertrand Matching, Claude Adam, Dominique Hasboun, Virginie Lambrecq, Vincent Navarro

## Abstract

Periventricular nodular heterotopia (PNH) is a malformation of cortical development that frequently causes drug-resistant epilepsy. The epileptogenicity of ectopic neurons in PNH as well as their role in generating interictal and ictal activity is still a matter of debate. We report the first *in vivo* microelectrode recording of heterotopic neurons in humans. Highly consistent interictal patterns (IPs) were identified within the nodules: 1) Periodic Discharges PLUS Fast activity (PD+F), Sporadic discharges PLUS Fast activity (SD+F), and 3) epileptic spikes (ES). Neuronal firing rates were significantly modulated during all IPs, suggesting that multiple IPs were generated by the same local neuronal populations. Furthermore, firing rates closely followed IP morphologies. Among the different IPs, SD+FA pattern was found only in the three nodules that were actively involved in seizure generation, but was never observed in the nodule that did not take part in ictal discharges. On the contrary, PD+F and ES were identified in all nodules. Units that were modulated during the IPs were also found to participate in seizures, increasing their firing rate at seizure onset and maintaining an elevated rate during the seizures. Together, nodules in PNH are highly epileptogenic, and show several IPs that provide promising pathognomonic signatures of PNH. Furthermore, our results show that PNH nodules may well initiate seizures.

**Highlights:**

- First *in vivo* microelectrode description of local epileptic activities in human PNH
- Recordings revealed multiple microscopic epileptic interictal patterns
- Firing rates of *all* detected units were significantly modulated during *all* interictal patterns
- Seizures recruited the same units that are involved in interictal activity

## Introduction

Periventricular nodular heterotopia (PNH) is one of the most common types of malformation of cortical development (MCD), resulting from an abnormal neuronal migration process wherein clusters of neurons form nodular masses of gray matter close to the walls of the lateral ventricles. Stereoelectroencephalography (SEEG) is often required to identify the seizure onset zone (SOZ) and delineate the region to be potentially removed (Battaglia et al., 2005; Tassi, 2004) or partially coagulated (Cossu et al., 2018). In other types of MCD, the underlying epileptogenic process can sometimes be inferred from patterns of interictal activity that are known to correlate with specific epileptogenic lesions (Perucca et al., 2014; Di Giacomo et al., 2019). For example, patients with type-IIb focal cortical dysplasia often show localized and continuous periodic spikes or poly-spikes (Gambardella et al., 1996; Tassi et al., 2012), while persistent and rhythmic fast-frequency discharges of very high amplitude (>100 µV) are typical of lissencephaly (Gastaut et al., 1987).

Recently, SEEG studies have suggested that stereotypical patterns could help in the identification of PNH. Tassi et al., (Mirandola et al., 2017; Tassi, 2004) found interictal pathologic activity that consisted of low voltage low frequency, interrupted by frequent high voltage spikes, and followed by positive waves. These spike and wave complexes were shown to originate only from nodules, and were very similar across patients and nodules. Pizzo et al, (Pizzo et al., 2017) also found that interictal activity in heterotopic cortex generally displays low amplitude activity and is further characterized by spikes and spike-and-wave complexes. Fast activity was often found as well, often superimposed on the slow wave following the spike.

Neurons within the ectopic tissue are highly disorganized (Dubeau et al., 1995; Thom et al., 2004). This limits spatial summation of electric currents measured by standard SEEG electrodes. Furthermore, it would make local field potentials (LFPs) highly susceptible from spread of electrical activity originating in surrounding tissue. Intracranial *micro*electrodes, due to their high impedance, record LFPs and action potentials from a much more focal region (≈140µm³), and are less dependent on the spatial summation of current flow (Buzsáki, 2006). This spatial selectivity allows the selective measurement of ectopic neuronal behavior by permitting the evaluation of interictal patterns recorded unequivocally within nodules. To the best of our knowledge, in-vivo recordings of single units have not yet been reported, and neither has interictal activity in PNH been measured with the spatial selectivity provided by microelectrodes. The investigation of the behavior of single units during interictal activity will therefore shed new light on the epileptogenicity of ectopic neurons in PNH, and their role in generating interictal activity.

The question whether nodules are typically part of the SOZ in PNH has been widely addressed in previous SEEG studies, showing that PNH involves complex, patient-specific epileptic networks that include nodules, as well as the overlying cortex and mesial temporal structures when adjacent to nodules (Aghakhani, 2005; Battaglia et al., 2005; Jacobs et al., 2009; Kothare et al., 1998; Mirandola et al., 2017; Pizzo et al., 2017; Tassi, 2004; Valton et al., 2008). The current study will further substantiate the involvement of ectopic neurons in seizures by investigating the involvement of ectopic neurons during both interictal activity and seizures.

Most of our knowledge of PNH on the cellular level comes from animal models. Preclinical PNH animal models (methylazoxymethanol rat) found abnormal bursting-like properties and prolonged tonic firing in PNH neurons (Colacitti et al., 1998), together with a lack of functional A-type Kv4.2 potassium channels (Castro et al., 2001). These findings suggest an increased excitability and decreased seizure thresholds in PNH neurons. In line with increased excitability in PNH, *in vitro* studies found a significant prolongation of GABAergic inhibitory postsynaptic potentials in heterotopic neurons (Calcagnotto et al., 2002; Kitaura et al., 2012) and reduced NR2A and NR2B subunit expression in methylazoxymethanol rat models, as well as in resected human PNH tissue (Finardi et al., 2006). These findings suggest a compensating drive against the otherwise hyper-excitability of the pathological network of PNH.

The current study is the first *in vivo* recording of extracellular action potentials in awake behaving human patients, allowing analyses of the firing behavior of ectopic neurons during interictal and ictal activities. Specifically, we: i) analyzed the activity of heterotopic neurons during inter-ictal patterns (IPs), ii) investigated whether interictal patterns previously described on macro electrodes are consistent with those found on the microscopic level, and iii) explored whether neurons that respond to interictal activity are also involved in seizures. For this purpose, IPs were described in microelectrode recordings from four PNH nodules in three patients. Single units were extracted using spike-sorting algorithms and their firing rate time-locked and correlated with the IPs. We expected to find i) consistent IPs within and between all four nodules, consistent with those described on macro electrodes, ii) hyper excitable neuronal firing during IPs, shown by a significant proportion of cells responding to interictal activity in a time-locked manner, and iii) that units that respond to inter-ictal activity are recruited during seizures.

## Materials and methods

### Patients

Three patients suffering from pharmacoresistant focal epilepsy due to PNH were included (see Table 1 and Supplementary Material). Patients received a full presurgical evaluation in the Epileptology Unit of the Pitié-Salpêtrière Hospital. All patients underwent 27-electrode long-term video-EEG recordings (Micromed System, Italy). The presurgical evaluation also included a structural 3T epilepsy-oriented brain MRI protocol, interictal 18FDG-PET and ictal (SISCOM) Tc-99m-HMPAO SPECT analysis. In Patient 1, MRI revealed a heterotopic nodule on the temporal horn of the right ventricle while Patient 2 presented with bilateral nodules predominantly in the posterior regions. In Patient 3, periventricular nodules were located in the posterior part of the right temporal lobe. In Patient 1 and 3, periventricular nodules were also associated with other anatomical abnormalities, including subcortical nodules (Patient 1 and 3), dysplastic overlying cortex (Patient 1) and polymicrogyric overlying temporal cortex (Patient 3). Data from Patient 1 were analyzed for 20.6 consecutive hours. Data from Patient 2, due to an abundance of interictal activity, were analyzed for 6 hours. In Patient 3, interictal activity was analyzed for 17.5 consecutive hours (Table 2).

**Table 1.** Clinical summary. of the presurgical evaluation. **IEDs**: Interictal Epileptiform discharges. **FSLA:** Focal seizures with loss of awareness. **FTBS:** Focal to bilateral seizures. **PNH:** Periventricular Nodular Heterotopia**. PMG:** Polymicrogyria. **SNH:** Subcortical Nodular Heterotopia. **CBZ**: Carbamazepine. **LCS:** Lacosamide. **LTG**: Lamotrigine. **ZNG**: Zonisamide.

| Age at 1st seizure (Sex) | Type | Epilepsy duration until sEEG | IEDs (scalp EEG) | Seizures (Scalp EEG) | SOZ defined by sEEG and % from the total number of seizures | MRI | PET | SPECT | Med. |
| --- | --- | --- | --- | --- | --- | --- | --- | --- | --- |
| 20 (M) | FSLA<br>FTBS | 11 yrs | Right anterior temporal regions | Right temporal region, spread to bilateral frontal regions | 89% the nodular lesion (n=40).<br>11% (n=5) postero-inferior cortical malformations close to the right hippocampal formation | PNH in the temporal horn of the right ventricle. Adjacent SNH and dysplastic overlying cortex | Hypermetabolism PNH | - | LTG<br>CBZ<br>LCS<br>CBZ |
| 25 (F) | FSLA<br>FTBS | 5 yrs | Left temporo-occipital region | Left posterior temporal region | 74 % (n=14) left PNH.<br>25% (n=5) diffuse origin | Bilateral posterior PNH.<br>No associated cortical abnormalities | No metabolic changes of the PNH | Hyperperfusion of the cortical structures adjacent to the nodules.<br>No evident metabolic modification in PNH | LTG |
| 14 (F) | FSLA<br>FTBS | 16 yrs | Right temporal or fronto-temporal region. Focal fast oscillations (20-24Hz) in the right temporal lobe | Right temporal region | 44% (n=20) nodules. 38% (n=17) right posterior parahippocampal region.<br>18% (n=8) diffuse origin | Right temporo-posterior PNH<br>Temporal SNH and PMG | Isometabolism of the PNH, PMG and adjacent cortices | Ictal hyperperfusion of PNH, PMG and adjacent and right temporo-polar cortices | LTG<br>ZNG<br>LCS |

**Table 2.** Localisation of micro and macro electrode contacts close to PNH.

|  |  | Macro contacts |  |  |  |  |  |  |  |
| --- | --- | --- | --- | --- | --- | --- | --- | --- | --- |
| N | Micro | 1 | 2 | 3 | 4 | 5 | 6 | 7 | 8 |
| 1 (R) | GM PNH | GM PNH | GM PNH<br>*lat* | GM STS | GM STS | GM STS | GM SMG | GM SMG | XC |
| 2 (R) | GM PNH | GM PNH | GM PNH<br>*lat* | WM | GM STG | GM STS<br>*T2* | GM STS<br>*T2* | GM *T2* | GM *T2* |
| 3 (L) | GM PNH | GM PNH<br>*im* | GM PNH<br>*im* | LTO | LTO *T3* | WM | WM | GM/WM<br>*T3* | GM *T3* |
| 4 (R) | GM<br>PNH) | GM PNH | GM PNH | WM | WM | GM STS | GM/WM<br>*T1* | GM *T1* | XC |

### SEEG recording

All intracranial SEEG procedures were performed according to clinical practice and the epilepsy features of the patients (see clinical supplemental materials S1 for clinical details). In addition to macroelectrodes, Behnke-Fried (Fried et al., 1999) type macro-micro electrodes (AdTech®) were inserted in nodules (Table 2). Signals from macro- and microelectrodes were continuously and synchronously recorded at 4kHz and 32kHz, respectively, using a hardware filter at 0.01Hz (Atlas Recording System; NeuraLynx, Tucson, AZ, USA). Post-implantation electrode locations were based on a pre-implantation 3T 3D-MRI, post-implant 1.5T 3D-MRI and CT scan, integrated using the Epiloc toolbox, an in-house developed plugin for the 3D-Slicer visualization software (Fedorov et al., 2012; Pérez-García et al., 2015). Patients gave a written informed consent (project C11-16 conducted by INSERM and approved by the local ethic committee, CPP Paris VI).

### Visual analysis of micro-LFP patterns

Pathological activities were visually identified on microelectrode LFP signals, according to traditional morphological characteristics used in clinical practice, then classified into different interictal patterns (IPs). Criteria for each pattern were then used for a complete manual scoring of continuous data recorded at the first day after implantation. The annotations were performed on the first day after the implantation, when the quality of microelectrode signal is typically highest. For each nodule, the annotations of the IPs were based on the LFP from the same microwire, selected for the clearest morphology, using a software developed in-house (MUSE). We describe our classification according to standard guidelines (Hirsch et al., 2013, 2021). Albeit designed for the critical care unit, such standard terminology improves clarity in communication, and to the best of our knowledge no other such standard exists. For periodic patterns, only sequences with at least 3 consecutive slow waves were considered. On a subset of these patterns, a detailed analysis of their periodicity was performed, by calculating the percentage of deflections that deviated less than 25% from the average period within each pattern (Hirsch et al., 2013, 2021).

### Seizure onset zone based on video-SEEG

The seizure onset zone (SOZ) of each patient was determined as part of the standard clinical procedure, involving visual annotations and analyses of the entire video-SEEG recordings by trained epileptologists (VF, CA & VN). In addition, for every seizure during the entire recording, the SOZ was scored as either nodular, non-nodular (cortical or hippocampal) or diffuse, resulting in a percentage of nodular seizures per patient. Representative examples of SEEG recording during seizures are reported in Supplementary Fig. 3.

### Time-locked & time-frequency analysis

Analyses were performed with a combination of FieldTrip (Oostenveld et al., 2011) and custom MATLAB scripts (The Mathworks Inc., Natick, Massachusetts). Manual annotations of each pattern, in each nodule, were temporally aligned by maximizing the cross-correlation between electrode time courses. Data was then down-sampled to 1 kHz. Average LFPs were extracted based on the aligned time-courses. Time–frequency representations (TFRs) were calculated using a Hanning taper applied to a sliding time window in steps of 5 ms. The time window was 200 ms for Epileptic Spikes, and 400 ms for all other (longer) patterns. Average power in 60 Hz - 200 Hz was expressed in percentage change relative, to the baseline-period that was set at −1s to −0.5 s for Epileptic Spikes, and −2 s to −1 s relative to pattern onset (annotation) for the other patterns.

### Spike sorting

All spikes occurring during manually annotated artefacted periods were ignored. After selecting electrodes that showed multi-unit activity (MUA), data were temporally whitened, and spikes were automatically detected at 6 (Nodule 1, 2 & 4) or 5.5 (Nodule 3) median absolute deviations of high-pass filtered (*>*300Hz) data. A combination of density-based clustering and template matching algorithms were used to automatically cluster the detected spikes (*Spyking Circus*, Yger et al., 2018). Clusters were visually merged when considered similar, based on the spike morphology, firing rate, amplitude and cross-correlation. Clusters were evaluated as to whether they reflected putative single-unit activity (SUA) or multi-unit activity, based on the inter-spike-interval (ISI), the percentage of refractory period violation (RPV = ISI *<* 2 ms) and spike morphology.

### Spike time analyses

Spike-times were epoched and time-locked according to the aligned annotations. Average spike rates were calculated continuously at 1000 Hz, using a Gaussian smoothing kernel of 10 ms for Epileptic Spikes, and 50 ms for the longer patterns to better capture the slower modulations of firing rate. Correlations were calculated between average time-locked spike rates of each unit, and the average time-locked LFP of every pattern. Finally, spike rates of each trial were binned into 100 bins for statistical analyses. To determine the resting behavior of detected units, data was first split into 10 second time periods, excluding time periods that overlapped with IED or seizure occurrence. The mean firing rate and Coefficient of Variation (Shinomoto et al., 2003) was calculated over the remaining time periods. Spike trains were time-locked to the manually annotated onset of every seizure.

### Statistical analysis

To control for multiple comparisons and non-normal distributions of firing-rates, we performed nonparametric cluster-based permutation tests to determine time-periods where firing rates changed significantly from baseline (Maris & Oostenveld, 2007). A threshold of *p<*0.01 (first-level t-test) was used to determine contiguous clusters, after which a threshold of *p<*0.05 (one-sided correction) determined whether the clusters (sum of t-values) could be explained under permutation (monte-carlo *n=*10.000). Due to the limited number of seizures, no frequentist statistics were performed on firing rates during seizures. However, raster plots were created, allowing qualitative observations on whether units involved in IPs were also recruited during seizure onset.

### Data availability statement

All scripts are made available on github: https://github.com/stephenwhitmarsh/EpiCode/projects/pnh. Anonymous data can be made available on reasonable request.

## Results

### Interictal epileptic LFP activities revealed by microelectrodes in PNH

Three interictal patterns (IPs) were consistently detected by microelectrodes in the nodules (Fig. 1 & Supplementary Fig. 1).

**Figure 1.**
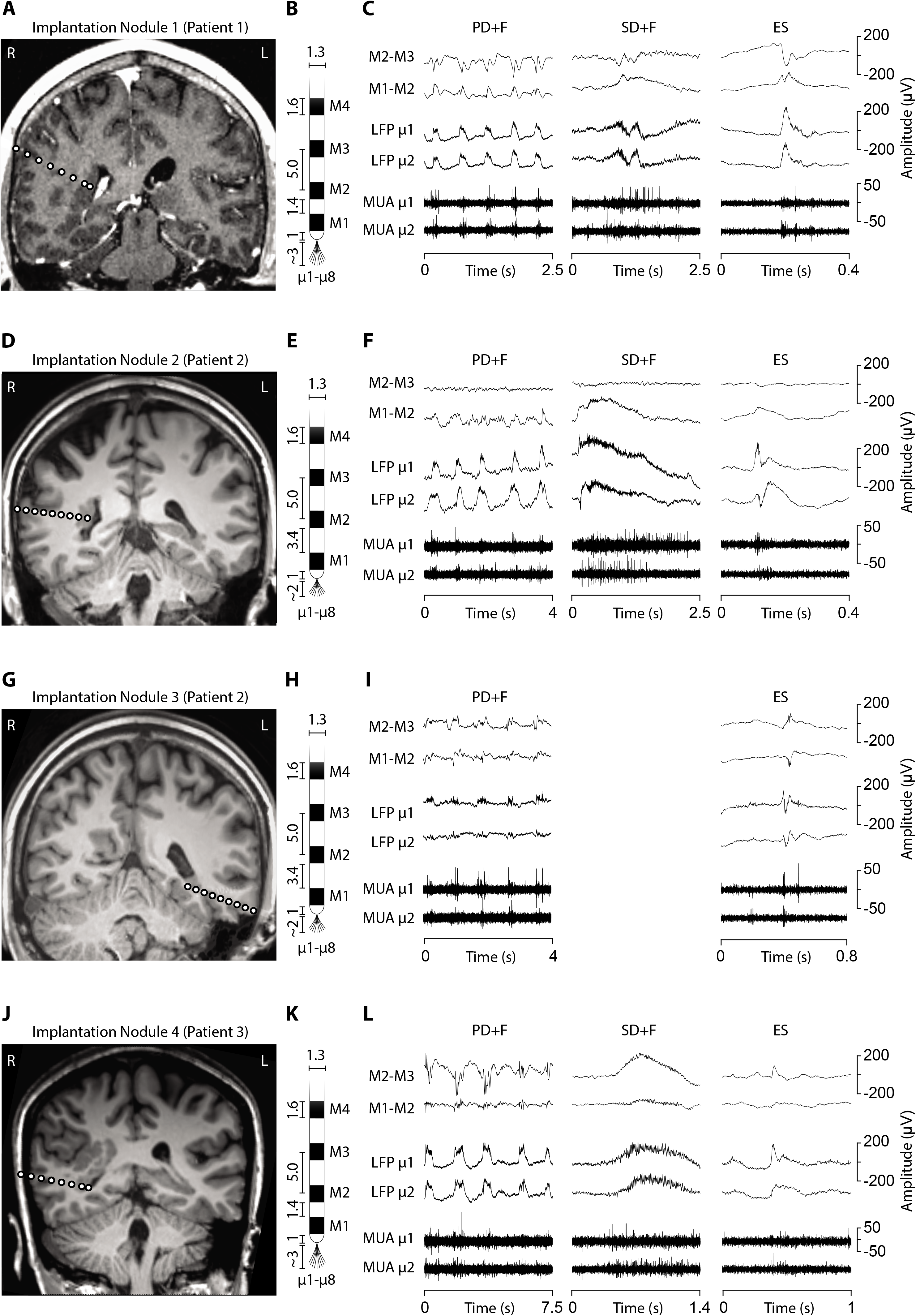
Electrode implantation into the PNH nodules and interictal LFP patterns. (A, D, G & J). Brain MRI showing the electrode trajectories, exploring the four analyzed nodules (A, Patient 1, Nodule 1; D & G, Patient 2, Nodule 2 & 3; J, Patient 3, Nodule 4). (B, E, H & K) Schematic representation of the macro-microelectrodes (M1-M4: macroelectrode contacts; μ1-μ8: microelectrodes). All the geometrical features of the electrodes are expressed in millimeters. Macroelectrode recordings (C, F, I & L) are shown in bipolar montage. Three LFP patterns were recorded: Periodic Discharge PLUS Fast activity (PD+F), Sporadic Discharge PLUS Fast activity (SD+F) and Epileptic Spikes (ES). These patterns were apparent on microelectrodes (LFP μ1 and μ2), and to a smaller degree also on macroelectrodes (M1-M2). LFP patterns were associated with MUA recorded on the microelectrodes.

#### Periodic Discharges PLUS Fast activity

We identified sequences of periodic slow waves, with superimposed low voltage fast activity. They were defined as Periodic Discharges PLUS Fast activity (PD+F) and were identified in all nodules (Fig. 1). These patterns were clearly detectable on microelectrodes, and to a smaller degree on the adjacent macroelectrode contacts (Fig. 1C, F & L, left panel). PD+F appeared as periodic bursts of fast activity on the closest macro contacts (M1-M2). In Nodule 3, Periodic Fast activity discharges (PF) consisted of periodic fast activity (PF; Fig. 1I, left panel). The PD+F patterns showed clear periodicity in the average LFP time course (Fig. 2). Consistent with the observation of low voltage fast activity in the single-trial time courses, the periodicity was also clearly visible in the frequency domain, with peak increases at 92 Hz, 135 Hz, 106 Hz and 81 Hz for Nodules 1, 2, 3 & 4, respectively (Fig. 2). A more detailed annotation within a subset of periodic patterns showed that the majority of deflections within a single train varied less than 25% of the mean interval between deflections (Table 3), conforming to the definition of *periodicity* (Hirsch et al., 2013, 2021).

**Figure 2.**
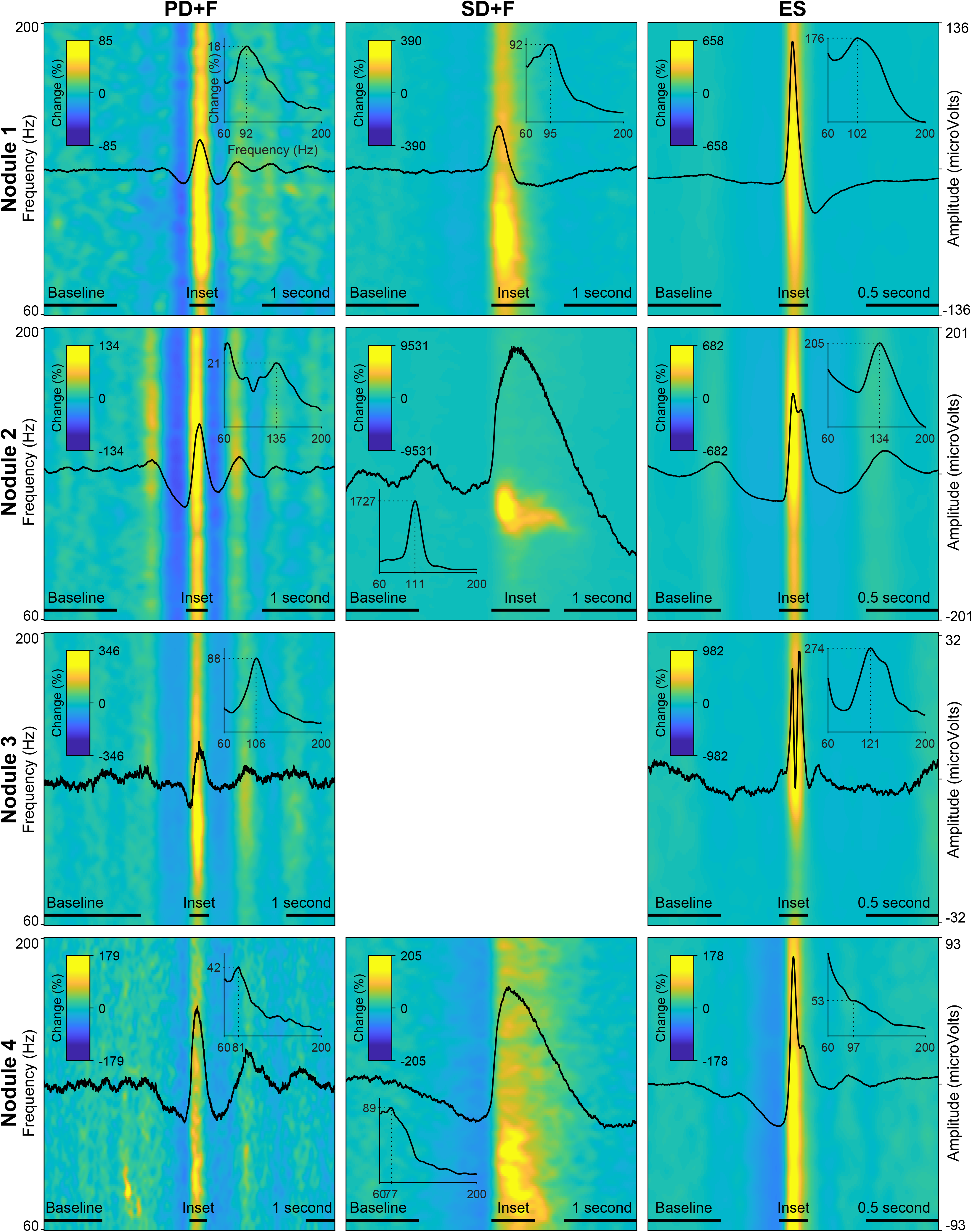
LFP and time-frequency spectra of the interictal patterns. Each row represents one nodule, with every panel/column an interictal pattern, indicated by the panel title. Amplitude of LFP are scaled per nodule, and displayed on the far-right y-axis of every row. Average time– frequency representations (60 Hz - 200Hz) are expressed in percentage change from the baseline-period, indicated by colorbar in every panel. Insets show average power spectra during peak activity of every pattern, indicated by horizontal bar above x-axis. The dotted lines in the inset show peak frequency and % power increased versus baseline. Note that due to the fact that these values are averaged over a time-segment, the percentage change is lower than indicated in the time-frequency plots.

**Table 3.** Descriptive statistics of (inter)ictal activity for each nodule. **N** = Nodule. **t** = Total duration of time analyzed (hrs.). **PD+F** = Periodic Discharges PLUS Fast activity. **SD+F** = Sporadic Discharge PLUS Fast activity. **ES** = Epileptic Spike. ***n*** = number of (inter)ictal events. λ = Mode of time between events (ms). xΔ**t** = Average time between periods (ms). ρΔ**t** = Standard deviation of time between periods (ms). **Periodicity** = ratio & percentage of periods that deviate less than 25% of the average time between periods.

| N | t | PD+F |  |  |  |  |  | SD+F |  | Epileptic Spike |  | Seizure |
| --- | --- | --- | --- | --- | --- | --- | --- | --- | --- | --- | --- | --- |
| | | *n* | $\lambda$ | $\bar{x} \Delta t$ | $\rho \Delta t$ | Periodicity | | *n* | $\lambda$ | *n* | $\lambda$ | *n* |
| 1 | 20,6 | 1485 | 291 | 511 | 18,7 | 124/150 | 83% | 950 | 302 | 3674 | 590 | 6 |
| 2 | 6,0 | 950 | 1445 | 574 | 20,6 | 93/117 | 80% | 89 | 4499 | 2685 | 494 | 0 |
| 3 | 6,0 | 518 | 1289 | 1074 | 17,6 | 116/128 | 91% |  |  | 392 | 373 | 0 |
| 4 | 17,5 | 231 | 4807 | 1723 | 19,6 | 62/72 | 86% | 507 | 1169 | 2504 | 361 | 8 |

#### Sporadic Discharges PLUS Fast Activity

We also identified large slow waves with superimposed and prolonged low voltage fast activity in Nodules 1, 3 & 4 (Figure 1C, F&L, middle panels). We termed this pattern Sporadic discharges PLUS Fast activity (SD+F). SD+F were also detectable on the macroelectrodes located in the nodules, despite smaller amplitudes (M1-M2 & M2-M3). These events were sporadic, with no tendency to further ictal organization. In Nodule 1, SD+F consisted of fast activity superimposed on a polyphasic slow deflection of 0.5-1s (Fig. 1C, middle panel), while in Nodule 2 and 4, fast activity occurred at the onset of a sharply appearing monophasic slow deflection of 2-4s (Fig. 1F and L, middle panel). Time-frequency analysis showed that, in Nodule 1, power increased at 96Hz for ≈ 0.5s (Fig. 2, top-middle panel). Nodule 2 showed a narrow-band increase of power at 111 Hz for at least one second (Fig. 2, center-middle panel). In Nodule 4, SD+F consisted of an increase of power in a broad frequency range, peaking at 77 Hz and lasting for 1-2 seconds (Fig. 2, lower-center panel).

#### Epileptic Spikes

Epileptic Spikes (ES) were identified in all four nodules on both the micro- and adjacent macro-electrodes contacts (Fig. 1 and Fig. 2). Nodule 1 showed isolated sharp monophasic waves (Fig. 1C, insert on the right). In Nodule 2 (Fig. 1F), spikes were generally followed, and often preceded, by a slow wave, which was also apparent in the average time-locked LFP. In Nodule 3 (Fig. 1I), spikes were characterized by a low amplitude di- or triphasic wave. In Nodule 4, spikes appeared as sharp monophasic waves often followed a slow wave (Fig. 1L, right panel). Low voltage fast activity was often superimposed on ES, with a mean peak at 102 Hz, 134 Hz, 121 Hz and 97 Hz, respectively. These ES were clearly visible also on macroelectrodes (Fig. 1C, F, I, L).

Patterns occurred frequently, often within seconds of each other. Table 3 reports the mode (λ) of each pattern, i.e., the most frequent time interval between events. The IPs correlated strongest with the LFPs of the closest macroelectrode contacts (M1-M2), after which the correlation reversed, showing that the IPs were located within the PNH (Supplementary Fig. 2).

### All neurons are recruited by interictal patterns

In addition to the LFP signals, microelectrodes recorded action potentials (multi-unit activity, MUA) in 24 microwires, located in periventricular nodules (Nodule 1: *n*=5, Nodule 2: *n*=7, Nodule 3: *n*=8, Nodule 4: *n*=4). Spike clustering resulted in 39 units of which 18 (55%) were classified as single units (Table 4). All units were shown to significantly modulate their firing rates in response to all IPs. Firing-rate increased up to 472% for PD+F, 10234% for SD+F, and 2138% for ES (Table 4). Most of the units also showed brief episodes of decreased firing rate surrounding the IPs, often by ≈100%, i.e., silence (Table 4 & Fig. 3). Especially during ES, firing rates strongly decreased in all units within ≈500 ms surrounding the discharge. Furthermore, correlations between firing rates and LFPs showed that the modulation of firing rates was highly consistent with the timing and shape of the IPs in all units (Table 4). A massive increase of firing rate was shown for the very short duration of the ES, while SD+F were associated with a more prolonged increase of firing rate (Fig. 3). During PD+F, firing rates showed regular and alternating periods of increases and decreases. Interestingly, two clusters (1 SUA & 1 MUA), significantly modulated their firing-rate inversely with respect to the IPs: they decreased their firing rate during interictal LFP activities (Table 4, Fig. 3: 4th row/neuron).

**Figure 3.**
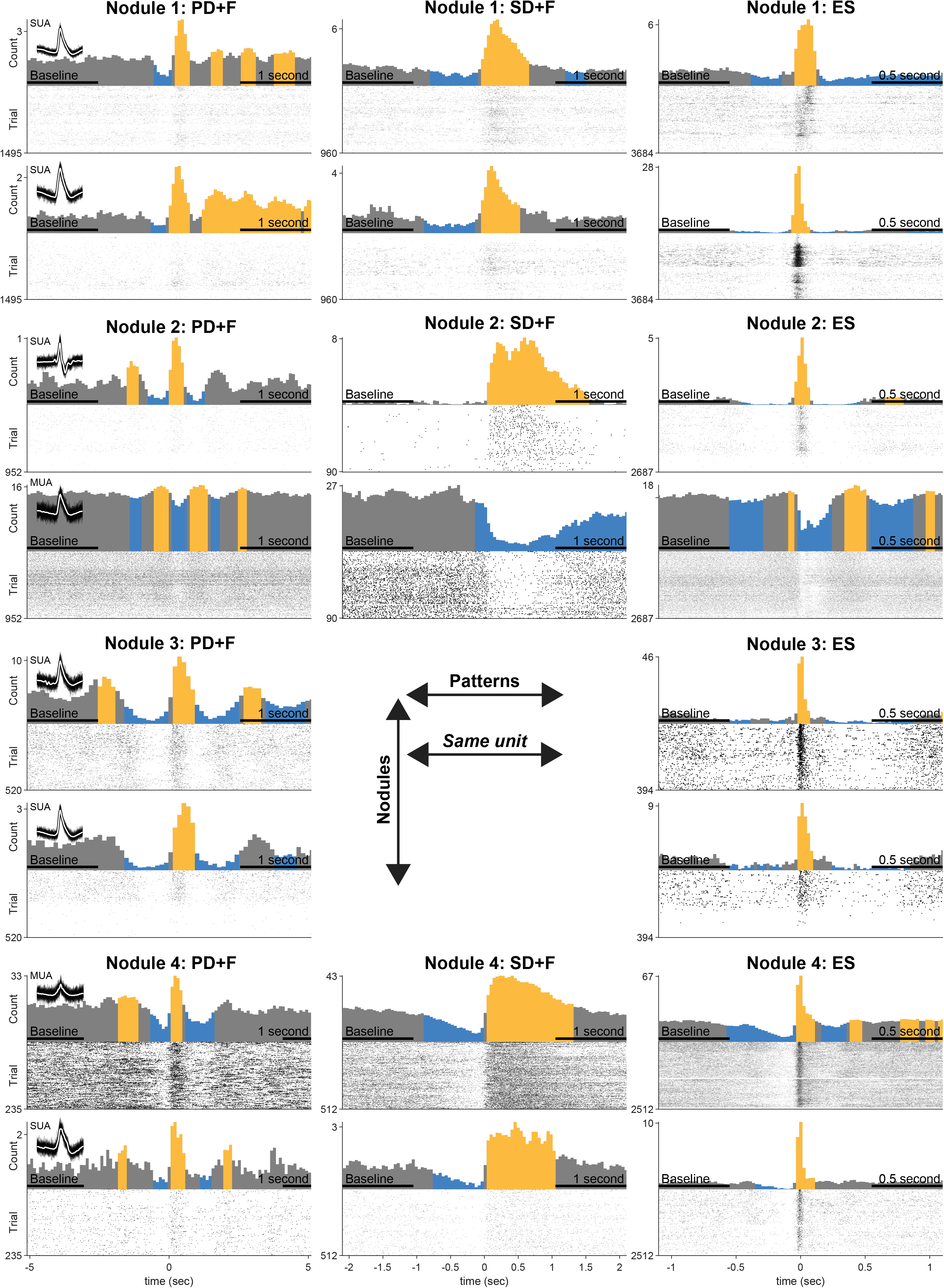
Statistical analysis of firing rate responses during IPs. Each row represents firing responses of one unit, shown with firing rate histogram centered on the onset of IP, above the respective raster plot. Colors indicate significant increases (yellow) and decreases (blue) compared to the average firing rate during baseline (indicated in x-axis). Two units are shown for every nodule (3 patients). Panel titles indicate interictal pattern. Note that each unit (row) shows significant modulation in every pattern. Inset shows 100 examples of action potential spikes, overlaid with their average spike waveform in white.

**Table 4.** Unit statistics. **U** = unit. **S/MU** = Single Unit Activity (SUA) or Multi Unit Activity (MUA). **RPV** = Refractory Period Violation (% > 2 ms). **PD+F** = Periodic Discharges PLUS Fast activity. **SD+F** = Sporadic Discharge PLUS Fast activity. **ES** = Epileptic Spike. Values under up (↑) and down (↓) arrows show the maximum percentage increase or decrease in significant clusters. ρ = pearson correlation between peri-stimulus time histogram and LFP. Negative correlations in bold font. Only significant values (**p** < 0.01) are shown. **FR**, **Amp** & **CV2** = Firing rate, average spike waveform amplitude & Coefficient of Variation, based on 10 second intervals without overlap with IEDs.

| U | Nodule | S/MU | RPV | PD+F |  |  | SD+F |  |  | ES |  |  | Baseline |  |  |
| --- | --- | --- | --- | --- | --- | --- | --- | --- | --- | --- | --- | --- | --- | --- | --- |
|  |  |  |  | ↑ | ↓ | ρ | ↑ | ↓ | ρ | ↑ | ↓ | ρ | FR | Amp | CV2 |
| 1 | 1 | SUA | 0,5 | 154 | 20 | 0,88 | 253 | 22 | 0,49 | 331 | 22 | 0,72 | 4,1 | 71,3 | 1,21 |
| 2 |  |  | 0,3 | 250 | *n.s.* | 0,59 | 472 | 52 | 0,74 | 426 | 61 | 0,24 | 15,2 | 115,0 | 1,19 |
| 3 |  |  | 1,7 | 124 | 16 | 0,89 | 227 | 21 | 0,54 | 261 | 17 | 0,76 | 4,1 | 49,1 | 1,18 |
| 4 |  |  | 0,7 | 154 | 38 | 0,77 | 257 | 33 | 0,63 | 1971 | 31 | 0,80 | 26,8 | 47,0 | 1,24 |
| 5 |  |  | 0,3 | 321 | 25 | 0,75 | 379 | 28 | 0,61 | 263 | 26 | 0,61 | 8,2 | 170,1 | 1,24 |
| 6 |  | MUA | 1,3 | 171 | 29 | 0,9 | 256 | 14 | 0,53 | 292 | 21 | 0,81 | 3,9 | 38,0 | 1,15 |
| 7 |  |  | 1,6 | 93 | 10 | 0,78 | 239 | 30 | 0,55 | 1098 | 17 | 0,91 | 3,8 | 28,3 | 1,17 |
| 8 |  |  | 1,9 | 352 | 53 | 0,72 | 454 | 52 | 0,59 | 441 | 56 | 0,30 | 25,5 | 67,5 | 1,18 |
| 9 |  |  | 3,0 | 121 | 15 | 0,90 | 231 | 12 | 0,53 | 366 | 14 | 0,84 | 5,3 | 30,9 | 1,16 |
| 10 | 2 | SUA | 0,3 | 192 | 35 | 0,51 | 460 | 49 | 0,27 | 597 | 21 | 0,50 | 2,6 | 55,4 | 1,02 |
| 11 |  |  | 0,2 | 199 | 40 | 0,74 | 3846 | 97 | 0,87 | 1325 | 48 | 0,75 | 2,3 | 69,6 | 1,27 |
| 12 |  |  | 1,1 | *n.s.* | 13 | 0,31 | *n.s.* | 36 | 0,28 | 79 | 15 | 0,71 | 6,5 | 29,8 | 1,04 |
| 13 |  |  | 0,1 | *n.s.* | 14 | **-0,26** | *n.s.* | 68 | **-0,39** | 36 | 20 | **-0,31** | 3,8 | 53,2 | 0,86 |
| 14 |  | MUA | 0,8 | 179 | 19 | 0,58 | 990 | 38 | 0,49 | 794 | 30 | 0,56 | 3,4 | 43,3 | 1,00 |
| 15 |  |  | 2,4 | 260 | 36 | 0,69 | 3193 | 99 | 0,72 | 962 | 41 | 0,75 | 5,5 | 29,5 | 1,15 |
| 16 |  |  | 0,8 | 247 | 40 | 0,51 | 10234 | *n.s.* | *n.s.* | 905 | 23 | 0,76 | 3,6 | 41,5 | 1,17 |
| 17 |  |  | 1,3 | 125 | 25 | 0,41 | *n.s.* | 86 | 0,08 | 363 | 29 | 0,67 | 41,8 | 65,9 | 1,09 |
| 18 |  |  | 4,0 | 161 | 19 | 0,70 | 1216 | 50 | 0,85 | 769 | 25 | 0,63 | 7,3 | 30,3 | 1,04 |
| 19 |  |  | 1,3 | 15 | 9 | **-0,79** | *n.s.* | 15 | **-0,62** | 19 | 16 | **-0,62** | 14,1 | 29,5 | 0,77 |
| 20 |  |  | 2,0 | 191 | 18 | 0,70 | 1480 | 51 | 0,68 | 621 | 15 | 0,83 | 3,7 | 30,7 | 1,17 |
| 21 |  |  | 2,4 | 255 | 31 | 0,80 | 4718 | *n.s.* | 0,88 | 1181 | 31 | 0,75 | 12,2 | 55,7 | 1,27 |
| 22 |  |  | 3,0 | 146 | 23 | 0,71 | 1381 | *n.s.* | 0,84 | 522 | 19 | 0,72 | 8,8 | 32,2 | 1,01 |
| 23 | 3 | SUA | 2,2 | 123 | 18 | 0,63 |  |  |  | 847 | 36 | 0,57 | 4,0 | 39,2 | 1,16 |
| 24 |  |  | 0,2 | 290 | 60 | 0,55 |  |  |  | *n.s.* | 100 | 0,53 | 5,8 | 80,2 | 1,24 |
| 25 |  |  | 0,3 | 426 | 54 | 0,59 |  |  |  | 2138 | 99 | 0,56 | 20,8 | 61,6 | 1,17 |
| 26 |  |  | 0,5 | 168 | 60 | 0,58 |  |  |  | 1087 | 66 | 0,45 | 2,2 | 33,7 | 1,18 |
| 27 |  |  | 0,7 | 143 | 35 | 0,72 |  |  |  | 484 | 61 | 0,75 | 2,2 | 71,6 | 1,19 |
| 28 |  |  | 0,5 | 368 | 35 | 0,69 |  |  |  | 819 | 76 | 0,62 | 1,5 | 40,4 | 1,29 |
| 29 |  |  | 0,1 | 343 | 37 | 0,51 |  |  |  | 1211 | 80 | 0,62 | 1,5 | 51,6 | 1,18 |
| 30 |  | MUA | 1,0 | 153 | 52 | 0,56 |  |  |  | 1398 | 41 | 0,51 | 2,2 | 33,5 | 1,18 |
| 31 |  |  | 0,6 | 117 | 46 | 0,54 |  |  |  | 655 | 58 | 0,52 | 1,4 | 31,2 | 1,12 |
| 32 |  |  | 2,1 | 176 | 51 | 0,70 |  |  |  | 625 | 45 | 0,53 | 3,6 | 19,7 | 1,25 |
| 33 |  |  | 0,3 | 472 | 48 | 0,58 |  |  |  | 1571 | 50 | 0,54 | 3,7 | 32,9 | 1,22 |
| 34 | 4 | SUA | 0,8 | *n.s.* | 33 | 0,45 | 241 | 32 | 0,81 | *n.s.* | 17 | 0,22 | 2,8 | 87,1 | 1,04 |
| 35 |  |  | 0,5 | 126 | 47 | 0,70 | 182 | 31 | 0,84 | 876 | 25 | 0,79 | 2,1 | 40,1 | 1,19 |
| 36 |  | MUA | 4,6 | 54 | 27 | 0,77 | 109 | 16 | 0,89 | 231 | 11 | 0,92 | 16,4 | 23,8 | 0,76 |
| 37 |  |  | 6,6 | 59 | 17 | 0,77 | 112 | 9 | 0,89 | 184 | 9 | 0,90 | 21,7 | 20,3 | 0,69 |
| 38 |  |  | 3,6 | 78 | 22 | 0,84 | 120 | 12 | 0,89 | 366 | 9 | 0,89 | 11,3 | 18,1 | 0,83 |
| 39 |  |  | 1,7 | *n.s.* | 54 | 0,51 | 155 | 42 | 0,82 | 323 | 56 | 0,73 | 2,3 | 59,0 | 1,12 |

### Nodules are involved in seizure initiation

The continuous two-week SEEG recordings allowed us to describe the complex ictogenesis of patients with PNH. In all patients, the seizures started predominantly in the PNH (from 42 to 90% of the total recorded seizures; see Table 1). In Patient 1, seizures from nodules were characterized by the appearance of a slow deflection overlapping with fast rhythms between 115 Hz and 160 Hz (Supplementary Fig. 3). Seizures started from the deep contacts of the two macroelectrodes, located in the posterior part of the PNH (Table 1), strongly implicating the PNH as the SOZ. In Patient 2, seizures were characterized by fast rhythms (above 270 Hz), starting in the contacts located within the anterior part of the left nodule, with a very fast involvement of the posterior part of homolateral PNH and the adjacent temporal cortex. The SOZ was thus represented by a complex network involving both the left PNH and the adjacent cortex. In Patient 2, the right nodule was not part of the SOZ but did eventually take part in later stages of seizure development. In Patient 3, focal seizures from the nodule started as a slow deflection with superimposed fast rhythms visible on the deepest contacts located in the most inferior and posterior part of the nodular lesions. Post-surgical validations of their SOZ were not available, because none of the included patients were selected for traditional surgical intervention due to the extent of the pathological lesions and the complexity of the epileptic networks.

### Units participating in the IPs are also recruited during seizures

In Patients 1 & 3, seizures were recorded during the periods in which MUA could be observed, allowing the observation of firing rate modulation during seizure initiation. In Patient 2, seizures occurred late during the continuous recordings, during a period where no MUA were present anymore. In Patient 1, all six seizures started in the nodule (Nodule 1). In Patient 3, six seizures started from the nodule (Nodule 4), one seizure had a diffuse onset, and one seizure could not be analyzed due to artifacts. In total, unit firing times were recovered during 12 seizures. The recovered units were the same as those analyzed for IPs. Although the relatively small number of observations precludes statistical tests, units were shown to increase their firing rate in time with seizure onset, and maintain an elevated rate for its duration (Fig. 4). This strongly suggests that units take part not only in different types of interictal activities, but are also recruited during seizure onset.

**Figure 4.**
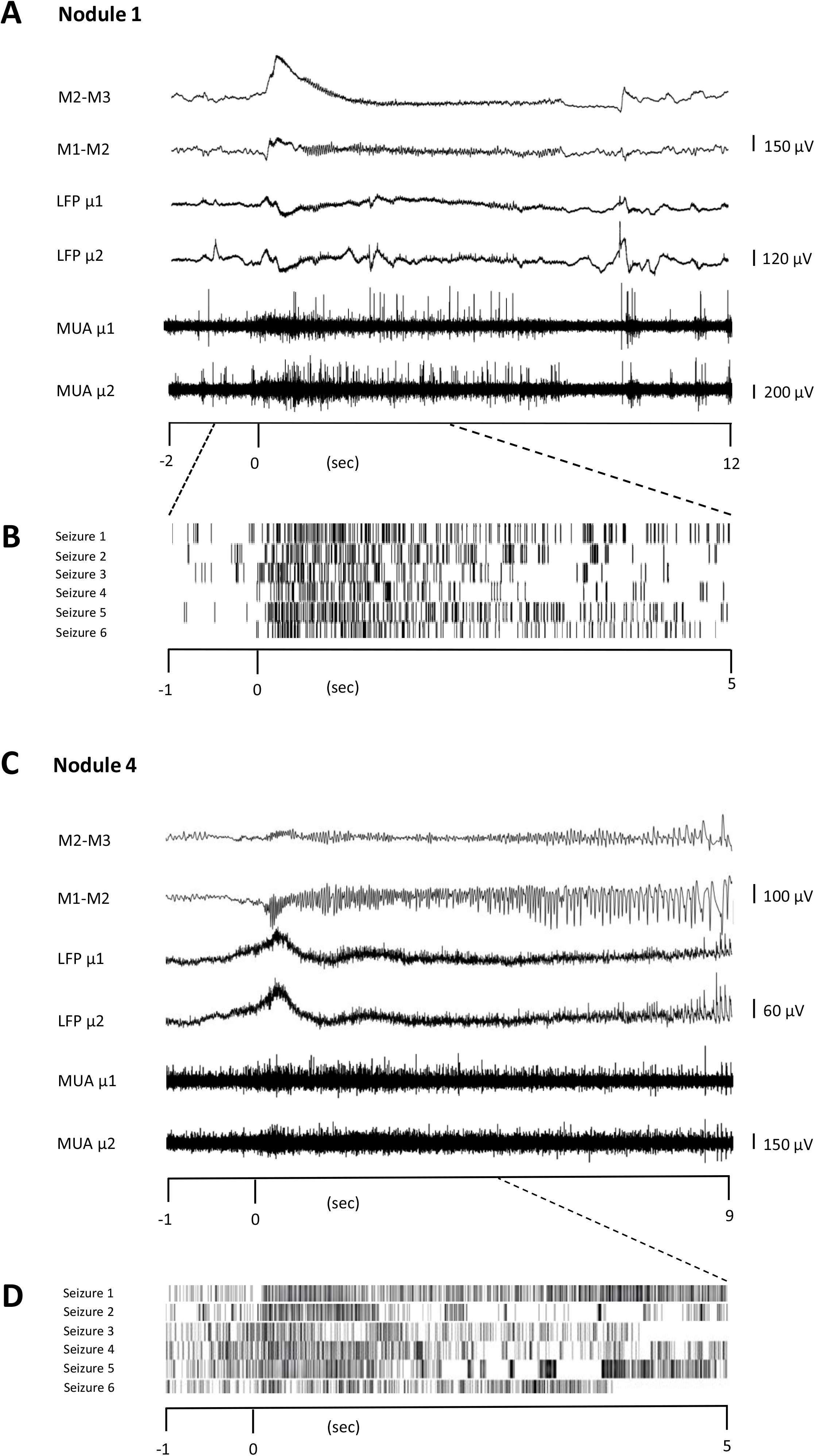
Unit firing behaviour during seizures. Two representative seizures originating from Nodule 1 (A) et Nodule 4 (C) respectively. The ictal activity from two adjacent contacts of the macroelectrode (M1-M2, M2-M3, bipolar montage) is reported together with that obtained from two different microelectrodes (µ1 and µ2). Microelectrode activity is then shown as LFP and MUA activity. TRaster plot, time-locked to seizure onset, for Nodule 1 (B) and Nodule 4 (D). In the raster plot, different shades of gray denote different units. Each line of the raster plot shows the unit’s behaviour during a seizure. The first line in the raster plot corresponds to the seizure shown above. Note that the units are the same as those analyzed for IEDs, here showing that they are also recruited during seizures.

## Discussion

Two novel interictal patterns (IPs) were identified in all four PNH nodules: Periodic Discharges PLUS Fast activity (PD+F) and Sporadic Discharges PLUS Fast Activity (SD+F). Epileptic Spikes (ES) were also shown to occur often and consistently in all four nodules. We showed that the IPs originated from the nodules and not from the adjacent cortex. First, the microelectrode locations were validated by neuroimaging to be located within the PNH. Second, due to their small size and high impedance, microelectrodes record highly localized activity in the order of 1 mm²3 (Buzsáki, 2004). Third, the LFPs only correlated with neighboring macro contacts that were also located within the PNH (Supplementary Fig. 2).

Microelectrode also allowed the recording of multiunit activity within PNH. We found that all spontaneously active neurons modulated their firing rates in accordance with the LFPs of all IPs. Such coincidence of neocortical neuronal discharge with IEDs recorded with surface EEG, i.e. “Positive” (Ishijima, 1972) or “involved” neurons (Prince & Futamachi, 1970), was suggested to be an indicator of the intensity of epileptic activity (Ishijima et al., 1975). Understanding the relationships between single unit activity and IEDs in the human brain has proven to be complex, however. Previous microelectrode studies found a strong heterogeneity in neuronal firing rates during cortical (Wyler et al., 1982), (Keller et al., 2010) as well during mesial temporal interictal discharges (Altafullah et al., 1986; Alvarado-Rojas et al., 2013), but none have, as of yet, reported on PNH. Some studies showed that only a subset of units increases its firing rate during the fast component of the epileptic spike (Altafullah et al., 1986; Isokawa-Akesson et al., 1989; Keller et al., 2010; Ulbert et al., 2004; Wyler et al., 1982). More consistent behavior was n found during the slow component of the ictal discharge, during which the firing rate is demising, independent of their cortical or limbic origin (Altafullah et al., 1986; Alvarado-Rojas et al., 2013; Keller et al., 2010; Ulbert et al., 2004). In the current study, both during the fast component of the epileptic spikes, as well as during the slow components, of both the epileptic spikes and novel periodic patterns, all the detected units significantly modulated their firing rates (Fig. 3), with the majority of units increasing during fast activity, and decreasing during slow activity (Table 4). Together, our results point to a highly pathological and reactive neuronal organization in PNH. This is in line with data obtained from rodent models, where PNH neurons showed aberrant cellular expression of ionic channels and thus cellular hyper-excitabity (Calcagnotto et al., 2002; Castro et al., 2001; Colacitti et al., 1998; Finardi et al., 2006). Indeed, heterotopic nodules show only a rudimentary laminar pattern and a lower degree of organization (Dubeau et al., 1995; Thom et al., 2004).

High Frequency Oscillations (HFOs) were shown to be superimposed onto all the three IPs. We found narrow-band increases of power in the high-gamma and ripple domain during SD+F (96, 111 & 77 Hz, in Nodule 1, 2 and 4, respectively), ES (102, 134, 121 & 97 Hz, in Nodule 1 to 4, respectively) and PD+F (92, 135, 106 & 81 Hz, in Nodule 1 to 4, respectively). Ripples (90-250 Hz) have been found in PNH nodules before, independent of whether the nodule was part of the SOZ (Ferrari-Marinho et al., 2015; Pizzo et al., 2017). In PNH patients, fast ripple (250-500 Hz) generation was identified in normotopic SOZ (Jacobs et al., 2009). Interestingly, in our findings, the SD+FA pattern was found only in the three nodules that were actively involved in seizure generation (Nodule 1, 2 and 4), but was never observed in the nodule that did not take part in ictal discharges (Nodule 3).

The role of PNH nodules in the initiation of seizures is still a matter of debate. Presurgical investigations in PNH patients suggest that periventricular heterotopia might best be considered as part of a complex pathological network that engages both nodules and cortical structures, resulting in seizures that might start in PNH, in the overlying cortex, or both (Kothare et al., 1998; Tassi, 2004; Aghakhani, 2005; Battaglia et al., 2005; Stefan et al., 2007; Valton et al., 2008; Jacobs et al., 2009; Mirandola et al., 2017; Pizzo et al., 2017). Studies on animal models of PNH showed that heterotopic nodules are reciprocally connected with both neocortical structures and the hippocampus (Chevassus-Au-Louis et al., 1998; Colacitti et al., 1998; Chevassus-au-Louis et al., 1999). Indeed, *in vitro* as well as *in vivo* electrophysiological recordings in MAM-rats also demonstrated that PNH can independently generate interictal epileptiform activity despite not being part of the SOZ (Tschuluun et al., 2005, 2011). In our data as well, all the explored nodules showed Epileptic Spikes, regardless of whether the nodules were the origin of seizures. As mentioned earlier, one interictal pattern (SD+FA) was found only in the three nodules that were actively involved in seizure generation (Nodule 1, 2 and 4), while it was never observed in the nodule that did not take part in ictal discharges (Nodule 3). This finding might serve as a potential signature of SOZ in PNH. Interestingly, our results indicate that the units that were active during interictal activity, were also involved in ictal activity, extending previous findings showing a varied response of neurons during seizures (Bower et al., 2012; Lambrecq et al., 2017; Schevon et al., 2012; Truccolo et al., 2011). Unfortunately, in the bilaterally implanted patient, MUAs were not present anymore at the time of seizures, so that a comparison between ictogenic nodule and not-ictogenic nodule could not be made.

To conclude, this study presents the first *in vivo* microscopic description of the behavior of ectopic neurons during several interictal epileptic patterns. Compared to other epileptic tissues, PNH presented as a relatively clearly defined structure that was easily targeted for implantation with micro-electrodes. Our study indicates consistent pathological, hyper-excitable activity within heterotopic nodules. On the level of the LFP, sporadic and periodic interictal patterns with superimposed fast activity might provide a pathognomonic signature of PNH. Future research will have to determine whether these IPs are specific to PNH or might be found in other types of cortical malformations as well.

## Acknowledgements

We would like to thank Pierre Pouget (ICM, Pitié-Salpêtrière hospital, Paris, France) and Fiona Francis (Institut du Fer à moulin, Paris, France) for the helpful discussions and comments on the manuscript. We are very grateful for the technical and nursing staff of the Epilepsy Unit, and would like to thank them for their help and expertise in caring for the patients and their recordings.

## Author contributions

**V.F., S.W.** & **V.N.** conceived and designed the study. **V.F.** identified and selected the different electrophysiological patterns and performed the clinico-anatomical analyses. **S.W.** designed and performed the quantitative time-signal analyses, spike-analyses and statistics. **V.F.**, **S.W.** & **V.N.** interpreted the data. **K.L.** collected, organized and preprocessed the physiological data. **P.Y.** wrote the software for spike sorting and provided advice on the analysis. **J-D.L.** wrote the software for manual annotations for our analyses. **V.F. & D.H**. planned the trajectories of the macro-microelectrodes and their anatomical targets. **B.M.** performed the surgical implantations. **C.A.**, **V.F., D.H.**, **V.L.** & **V.N.** performed the clinical evaluations. **V.F.** & **S.W.** wrote the initial draft of the manuscript. **V.F.**, **S.W., V.L. & V.N.** provided critical revisions of the manuscript. All authors contributed to the final version of the manuscript

## Supplementary Data Description

**Supplementary Material** Clinical data.

**Supplementary Figure 1.** *Example of multichannel data showing three patterns within a single time period*.

**Supplementary Figure 2.** *Correlation LFP between micro and macro contacts.* For each Nodule and each pattern, the microelectrode LFP is correlated with each macro-contact of the same shaft (1=closest to microelectrode, 8 = farthest from microelectrode). Macro-contacts were bi-polar referenced. Numbers and size of markers denote positive (yellow) and negative (blue) correlation values. Note the largests correlation with the closest macro contact, as well as the flip of correlation values in the next contact, showing that the patterns were highly localized to the tip of the shaft.

**Supplementary Figure 3.** *Representative SEEG recordings obtained from the three patients during the clinical evaluation*

## Bibliography

Aghakhani, Y. (2005). The role of periventricular nodular heterotopia in epileptogenesis. Brain, 128(3), 641–651. https://doi.org/10.1093/brain/awh388

Altafullah, I., Halgren, E., Stapleton, J. M., & Crandall, P. H. (1986). Interictal spike-wave complexes in the human medial temporal lobe: Typical topography and comparisons with cognitive potentials. Electroencephalography and Clinical Neurophysiology, 63(6), 503– 516. https://doi.org/10.1016/0013-4694(86)90138-0

Alvarado-Rojas, C., Lehongre, K., Bagdasaryan, J., Bragin, A., Staba, R., Engel, J., Navarro, V., & Le Van Quyen, M. (2013). Single-unit activities during epileptic discharges in the human hippocampal formation. Frontiers in Computational Neuroscience, 7, 140. https://doi.org/10.3389/fncom.2013.00140

Battaglia, G., Franceschetti, S., Chiapparini, L., Freri, E., Bassanini, S., Giavazzi, A., Finardi, A., Taroni, F., & Granata, T. (2005). Electroencephalographic Recordings of Focal Seizures in Patients Affected by Periventricular Nodular Heterotopia: Role of the Heterotopic Nodules in the Genesis of Epileptic Discharges. Journal of Child Neurology, 20(4), 369– 377. https://doi.org/10.1177/08830738050200041701

Bower, M. R., Stead, M., Meyer, F. B., Marsh, W. R., & Worrell, G. A. (2012). Spatiotemporal neuronal correlates of seizure generation in focal epilepsy: Neuronal Correlates Outside the Focus. Epilepsia, 53(5), 807–816. https://doi.org/10.1111/j.1528-1167.2012.03417.x

Buzsáki, G. (2004). Large-scale recording of neuronal ensembles. Nature Neuroscience, 7(5), 446–451. https://doi.org/10.1038/nn1233

Buzsáki, G. (2006). Rhythms of the brain. Oxford Univ. Press.

Calcagnotto, M. E., Paredes, M. F., & Baraban, S. C. (2002). Heterotopic neurons with altered inhibitory synaptic function in an animal model of malformation-associated epilepsy. The Journal of Neuroscience: The Official Journal of the Society for Neuroscience, 22(17), 7596–7605.

Castro, P. A., Cooper, E. C., Lowenstein, D. H., & Baraban, S. C. (2001). Hippocampal heterotopia lack functional Kv4.2 potassium channels in the methylazoxymethanol model of cortical malformations and epilepsy. The Journal of Neuroscience: The Official Journal of the Society for Neuroscience, 21(17), 6626–6634.

Chevassus-au-Louis, N., Baraban, S. C., Gaïarsa, J. L., & Ben-Ari, Y. (1999). Cortical malformations and epilepsy: New insights from animal models. Epilepsia, 40(7), 811– 821. https://doi.org/10.1111/j.1528-1157.1999.tb00786.x

Chevassus-Au-Louis, N., Congar, P., Represa, A., Ben-Ari, Y., & Gaïarsa, J. L. (1998). Neuronal migration disorders: Heterotopic neocortical neurons in CA1 provide a bridge between the hippocampus and the neocortex. Proceedings of the National Academy of Sciences of the United States of America, 95(17), 10263–10268. https://doi.org/10.1073/pnas.95.17.10263

Colacitti, C., Sancini, G., Franceschetti, S., Cattabeni, F., Avanzini, G., Spreafico, R., Di Luca, M., & Battaglia, G. (1998). Altered connections between neocortical and heterotopic areas in methylazoxymethanol-treated rat. Epilepsy Research, 32(1–2), 49–62. https://doi.org/10.1016/s0920-1211(98)00039-4

Cossu, M., Mirandola, L., & Tassi, L. (2018). RF-ablation in periventricular heterotopia-related epilepsy. Epilepsy Research, 142, 121–125. https://doi.org/10.1016/j.eplepsyres.2017.07.001

Di Giacomo, R., Uribe-San-Martin, R., Mai, R., Francione, S., Nobili, L., Sartori, I., Gozzo, F., Pelliccia, V., Onofrj, M., Lo Russo, G., de Curtis, M., & Tassi, L. (2019). Stereo-EEG ictal/interictal patterns and underlying pathologies. Seizure, 72, 54–60. https://doi.org/10.1016/j.seizure.2019.10.001

Dubeau, F., Tampieri, D., Lee, N., Andermann, E., Carpenter, S., Leblanc, R., Olivier, A., Radtke, R., Villemure, J. G., & Andermann, F. (1995). Periventricular and subcortical nodular heterotopia A study of 33 patients. Brain, 118(5), 1273–1287. https://doi.org/10.1093/brain/118.5.1273

Fedorov, A., Beichel, R., Kalpathy-Cramer, J., Finet, J., Fillion-Robin, J.-C., Pujol, S., Bauer, C., Jennings, D., Fennessy, F., Sonka, M., Buatti, J., Aylward, S., Miller, J. V., Pieper, S., & Kikinis, R. (2012). 3D Slicer as an image computing platform for the Quantitative Imaging Network. Magnetic Resonance Imaging, 30(9), 1323–1341. https://doi.org/10.1016/j.mri.2012.05.001

Ferrari-Marinho, T., Perucca, P., Mok, K., Olivier, A., Hall, J., Dubeau, F., & Gotman, J. (2015). Pathologic substrates of focal epilepsy influence the generation of high-frequency oscillations. Epilepsia, 56(4), 592–598. https://doi.org/10.1111/epi.12940

Finardi, A., Gardoni, F., Bassanini, S., Lasio, G., Cossu, M., Tassi, L., Caccia, C., Taroni, F., LoRusso, G., Di Luca, M., & Battaglia, G. (2006). NMDA receptor composition differs among anatomically diverse malformations of cortical development. Journal of Neuropathology and Experimental Neurology, 65(9), 883–893. https://doi.org/10.1097/01.jnen.0000235117.67558.6d

Fried, I., Wilson, C. L., Maidment, N. T., Engel, J., Behnke, E., Fields, T. A., MacDonald, K. A., Morrow, J. W., & Ackerson, L. (1999). Cerebral microdialysis combined with single-neuron and electroencephalographic recording in neurosurgical patients. Technical note. Journal of Neurosurgery, 91(4), 697–705. https://doi.org/10.3171/jns.1999.91.4.0697

Gambardella, A., Palmini, A., Andermann, F., Dubeau, F., Da Costa, J. C., Felipe Quesney, L., Andermann, E., & Olivier, A. (1996). Usefulness of focal rhythmic discharges on scalp EEG of patients with focal cortical dysplasia and intractable epilepsy. Electroencephalography and Clinical Neurophysiology, 98(4), 243–249. https://doi.org/10.1016/0013-4694(95)00266-9

Gastaut, H., Pinsard, N., Raybaud, C., Aicardi, J., & Zifkin, B. (1987). Lissencephaly (agyria-pachygyria): Clinical findings and serial EEG studies. Developmental Medicine and Child Neurology, 29(2), 167–180. https://doi.org/10.1111/j.1469-8749.1987.tb02132.x

Hirsch, L. J., Fong, M. W. K., Leitinger, M., LaRoche, S. M., Beniczky, S., Abend, N. S., Lee, J. W., Wusthoff, C. J., Hahn, C. D., Westover, M. B., Gerard, E. E., Herman, S. T., Haider, H. A., Osman, G., Rodriguez-Ruiz, A., Maciel, C. B., Gilmore, E. J., Fernandez, A., Rosenthal, E. S., … Gaspard, N. (2021). American Clinical Neurophysiology Society’s Standardized Critical Care EEG Terminology: 2021 Version. Journal of Clinical Neurophysiology, 38(1), 1–29. https://doi.org/10.1097/WNP.0000000000000806

Hirsch, L. J., LaRoche, S. M., Gaspard, N., Gerard, E., Svoronos, A., Herman, S. T., Mani, R., Arif, H., Jette, N., Minazad, Y., Kerrigan, J. F., Vespa, P., Hantus, S., Claassen, J., Young, G. B., So, E., Kaplan, P. W., Nuwer, M. R., Fountain, N. B., & Drislane, F. W. (2013). American Clinical Neurophysiology Society’s Standardized Critical Care EEG Terminology: 2012 version. Journal of Clinical Neurophysiology, 30(1), 27.

Ishijima, B. (1972). Unitary analysis of epileptic activity in acute and chronic foci and related cortex of cat and monkey. Epilepsia, 13(4), 561–581. https://doi.org/10.1111/j.1528-1157.1972.tb04393.x

Ishijima, B., Hori, T., Yoshimasa, N., Fukushima, T., Hirakawa, K., & Sekino, H. (1975). Neuronal activities in human epileptic foci and surrounding areas. Electroencephalography and Clinical Neurophysiology, 39(6), 643–650. https://doi.org/10.1016/0013-4694(75)90077-2

Isokawa-Akesson, M., Wilson, C. L., & Babb, T. L. (1989). Inhibition in synchronously firing human hippocampal neurons. Epilepsy Research, 3(3), 236–247. https://doi.org/10.1016/0920-1211(89)90030-2

Jacobs, J., Levan, P., Châtillon, C.-E., Olivier, A., Dubeau, F., & Gotman, J. (2009). High frequency oscillations in intracranial EEGs mark epileptogenicity rather than lesion type. Brain: A Journal of Neurology, 132(Pt 4), 1022–1037. https://doi.org/10.1093/brain/awn351

Keller, C. J., Truccolo, W., Gale, J. T., Eskandar, E., Thesen, T., Carlson, C., Devinsky, O., Kuzniecky, R., Doyle, W. K., Madsen, J. R., Schomer, D. L., Mehta, A. D., Brown, E. N., Hochberg, L. R., Ulbert, I., Halgren, E., & Cash, S. S. (2010). Heterogeneous neuronal firing patterns during interictal epileptiform discharges in the human cortex. Brain, 133(6), 1668–1681. https://doi.org/10.1093/brain/awq112

Kitaura, H., Oishi, M., Takei, N., Fu, Y.-J., Hiraishi, T., Fukuda, M., Takahashi, H., Shibuki, K., Fujii, Y., & Kakita, A. (2012). Periventricular nodular heterotopia functionally couples with the overlying hippocampus: Optical Imaging of PVNH. Epilepsia, 53(7), e127– e131. https://doi.org/10.1111/j.1528-1167.2012.03509.x

Kothare, S. V., VanLandingham, K., Armon, C., Luther, J. S., Friedman, A., & Radtke, R. A. (1998). Seizure onset from periventricular nodular heterotopias: Depth-electrode study. Neurology, 51(6), 1723–1727. https://doi.org/10.1212/WNL.51.6.1723

Lambrecq, V., Lehongre, K., Adam, C., Frazzini, V., Mathon, B., Clemenceau, S., Hasboun, D., Charpier, S., Baulac, M., Navarro, V., & Le Van Quyen, M. (2017). Single-unit activities during the transition to seizures in deep mesial structures: Seizures and Single-Unit Activities. Annals of Neurology, 82(6), 1022–1028. https://doi.org/10.1002/ana.25111

Maris, E., & Oostenveld, R. (2007). Nonparametric statistical testing of EEG- and MEG-data. Journal of Neuroscience Methods, 164(1), 177–190. https://doi.org/10.1016/j.jneumeth.2007.03.024

Mirandola, L., Mai, R. F., Francione, S., Pelliccia, V., Gozzo, F., Sartori, I., Nobili, L., Cardinale, F., Cossu, M., Meletti, S., & Tassi, L. (2017). Stereo-EEG: Diagnostic and therapeutic tool for periventricular nodular heterotopia epilepsies. Epilepsia, 58(11), 1962–1971. https://doi.org/10.1111/epi.13895

Oostenveld, R., Fries, P., Maris, E., & Schoffelen, J.-M. (2011). FieldTrip: Open Source Software for Advanced Analysis of MEG, EEG, and Invasive Electrophysiological Data. Computational Intelligence and Neuroscience, 2011, 1–9. https://doi.org/10.1155/2011/156869

Pérez-García, F., Lehongre, K., Bardinet, E., Jannin, P., Navarro, V., Hasboun, D., & Fernandez-Vidal, S. (2015). Automatic Segmentation Of Depth Electrodes Implanted In Epileptic Patients: A Modular Tool Adaptable To Multicentric Protocols. Epilepsia, 56, 227.

Perucca, P., Dubeau, F., & Gotman, J. (2014). Intracranial electroencephalographic seizure-onset patterns: Effect of underlying pathology. Brain, 137(1), 183–196. https://doi.org/10.1093/brain/awt299

Pizzo, F., Roehri, N., Catenoix, H., Medina, S., McGonigal, A., Giusiano, B., Carron, R., Scavarda, D., Ostrowsky, K., Lepine, A., Boulogne, S., Scholly, J., Hirsch, E., Rheims, S., Bénar, C.-G., & Bartolomei, F. (2017). Epileptogenic networks in nodular heterotopia: A stereoelectroencephalography study. Epilepsia, 58(12), 2112–2123. https://doi.org/10.1111/epi.13919

Prince, D. A., & Futamachi, K. J. (1970). Intracellular recordings from chronic epileptogenic foci in the monkey. Electroencephalography and Clinical Neurophysiology, 29(5), 496– 510. https://doi.org/10.1016/0013-4694(70)90066-0

Schevon, C. A., Weiss, S. A., McKhann, G., Goodman, R. R., Yuste, R., Emerson, R. G., & Trevelyan, A. J. (2012). Evidence of an inhibitory restraint of seizure activity in humans. Nature Communications, 3, 1060. https://doi.org/10.1038/ncomms2056

Shinomoto, S., Shima, K., & Tanji, J. (2003). Differences in spiking patterns among cortical neurons. Neural Computation, 15(12), 2823–2842. https://doi.org/10.1162/089976603322518759

Stefan, H., Nimsky, C., Scheler, G., Rampp, S., Hopfengärtner, R., Hammen, T., Dörfler, A., Blümcke, I., & Romstöck, J. (2007). Periventricular nodular heterotopia: A challenge for epilepsy surgery. Seizure, 16(1), 81–86. https://doi.org/10.1016/j.seizure.2006.10.004

Tassi, L. (2004). Electroclinical, MRI and neuropathological study of 10 patients with nodular heterotopia, with surgical outcomes. Brain, 128(2), 321–337. https://doi.org/10.1093/brain/awh357

Tassi, L., Garbelli, R., Colombo, N., Bramerio, M., Russo, G. L., Mai, R., Deleo, F., Francione, S., Nobili, L., & Spreafico, R. (2012). Electroclinical, MRI and surgical outcomes in 100 epileptic patients with type II FCD. Epileptic Disorders: International Epilepsy Journal with Videotape, 14(3), 257–266. https://doi.org/10.1684/epd.2012.0525

Thom, M., Martinian, L., Parnavelas, J. G., & Sisodiya, S. M. (2004). Distribution of Cortical Interneurons in Grey Matter Heterotopia in Patients with Epilepsy. Epilepsia, 45(8), 916– 923. https://doi.org/10.1111/j.0013-9580.2004.46603.x

Truccolo, W., Donoghue, J. A., Hochberg, L. R., Eskandar, E. N., Madsen, J. R., Anderson, W. S., Brown, E. N., Halgren, E., & Cash, S. S. (2011). Single-neuron dynamics in human focal epilepsy. Nature Neuroscience, 14(5), 635–641. https://doi.org/10.1038/nn.2782

Tschuluun, N., Jürgen Wenzel, H., Doisy, E. T., & Schwartzkroin, P. A. (2011). Initiation of epileptiform activity in a rat model of periventricular nodular heterotopia: *Epileptiform Activity in Rat PNH*. Epilepsia, 52(12), 2304–2314. https://doi.org/10.1111/j.1528-1167.2011.03264.x

Tschuluun, N., Wenzel, J. H., Katleba, K., & Schwartzkroin, P. A. (2005). Initiation and spread of epileptiform discharges in the methylazoxymethanol acetate rat model of cortical dysplasia: Functional and structural connectivity between CA1 heterotopia and hippocampus/neocortex. Neuroscience, 133(1), 327–342. https://doi.org/10.1016/j.neuroscience.2005.02.009

Ulbert, I., Heit, G., Madsen, J., Karmos, G., & Halgren, E. (2004). Laminar analysis of human neocortical interictal spike generation and propagation: Current source density and multiunit analysis in vivo. Epilepsia, 45 *Suppl 4*, 48–56. https://doi.org/10.1111/j.0013-9580.2004.04011.x

Valton, L., Guye, M., McGonigal, A., Marquis, P., Wendling, F., Régis, J., Chauvel, P., & Bartolomei, F. (2008). Functional interactions in brain networks underlying epileptic seizures in bilateral diffuse periventricular heterotopia. Clinical Neurophysiology, 119(1), 212–223. https://doi.org/bartho

Wyler, A. R., Ojemann, G. A., & Ward, A. A. (1982). Neurons in human epileptic cortex: Correlation between unit and EEG activity. Annals of Neurology, 11(3), 301–308. https://doi.org/10.1002/ana.410110311

Yger, P., Spampinato, G. L., Esposito, E., Lefebvre, B., Deny, S., Gardella, C., Stimberg, M., Jetter, F., Zeck, G., Picaud, S., Duebel, J., & Marre, O. (2018). A spike sorting toolbox for up to thousands of electrodes validated with ground truth recordings in vitro and in vivo. ELife, 7. https://doi.org/10.7554/eLife.34518

