## Supplementary figures and images for "Human periventricular nodular heterotopia shows several interictal epileptic patterns, associated with hyperexcitability of neuronal firing"

### Supplementary Fig. 1

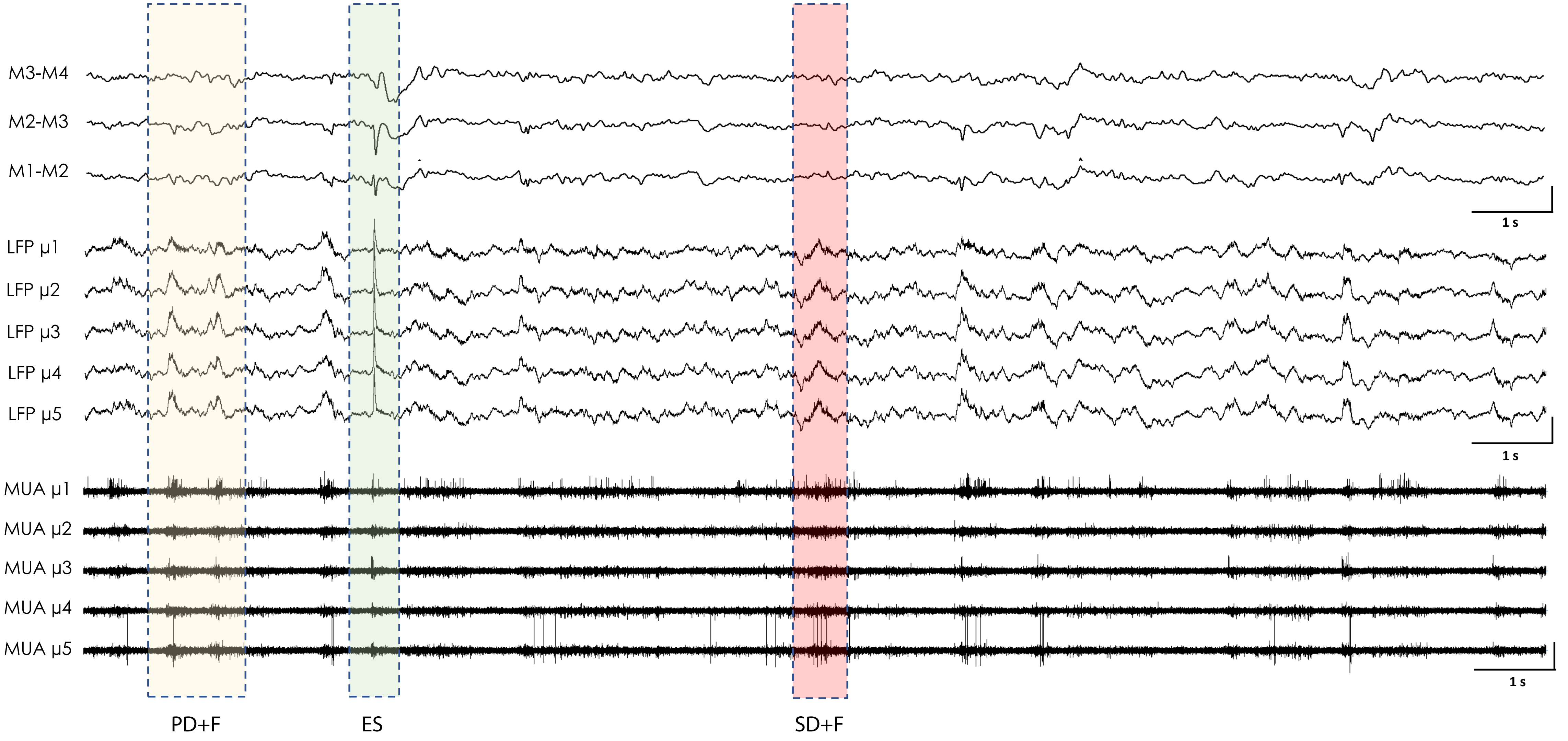

### Supplementary Fig. 2

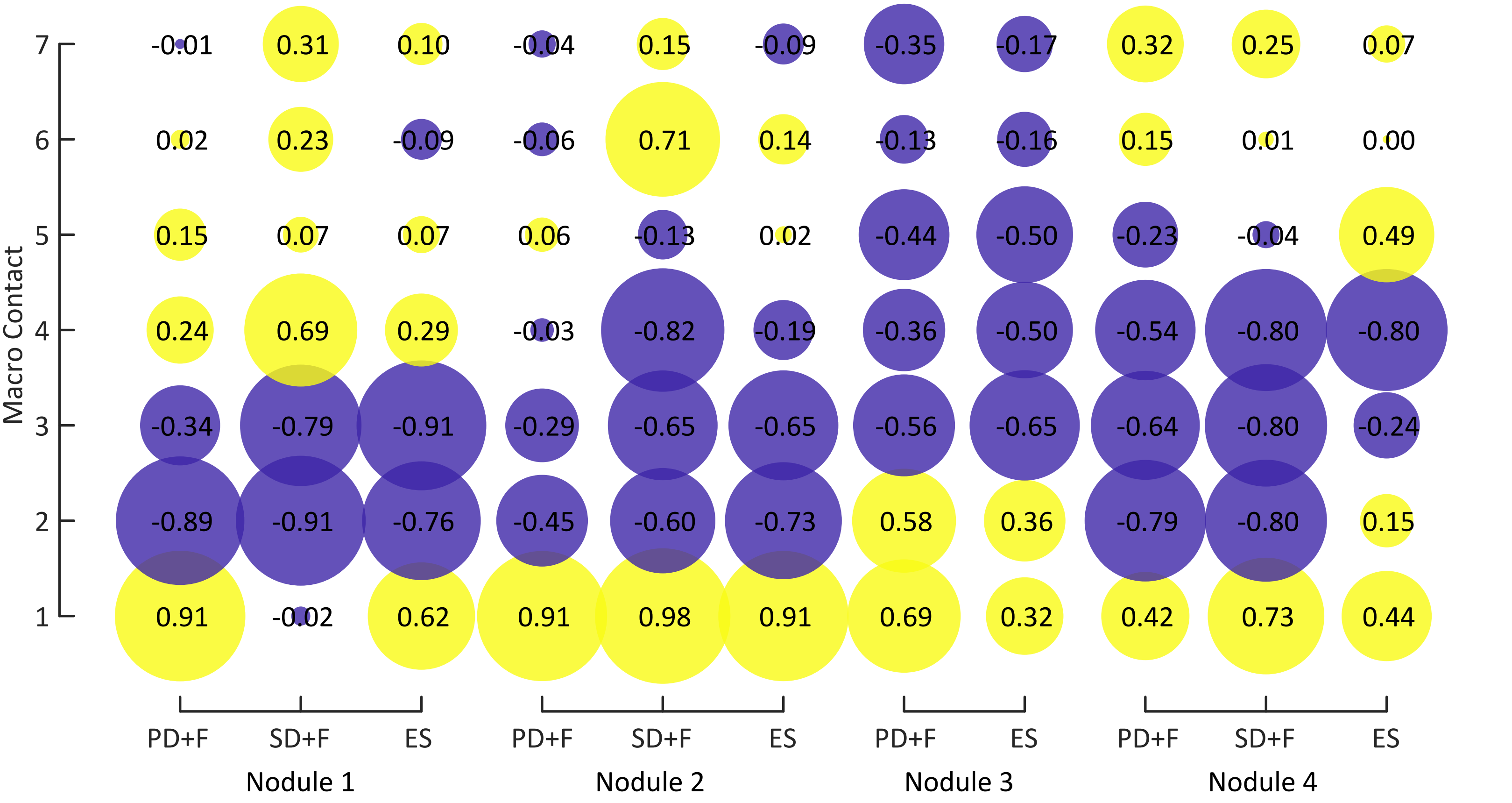
