## Supplementary Fig. 3 for "Human periventricular nodular heterotopia shows several interictal epileptic patterns, associated with hyperexcitability of neuronal firing"

Supplementary Fig. 3A. Patient 1

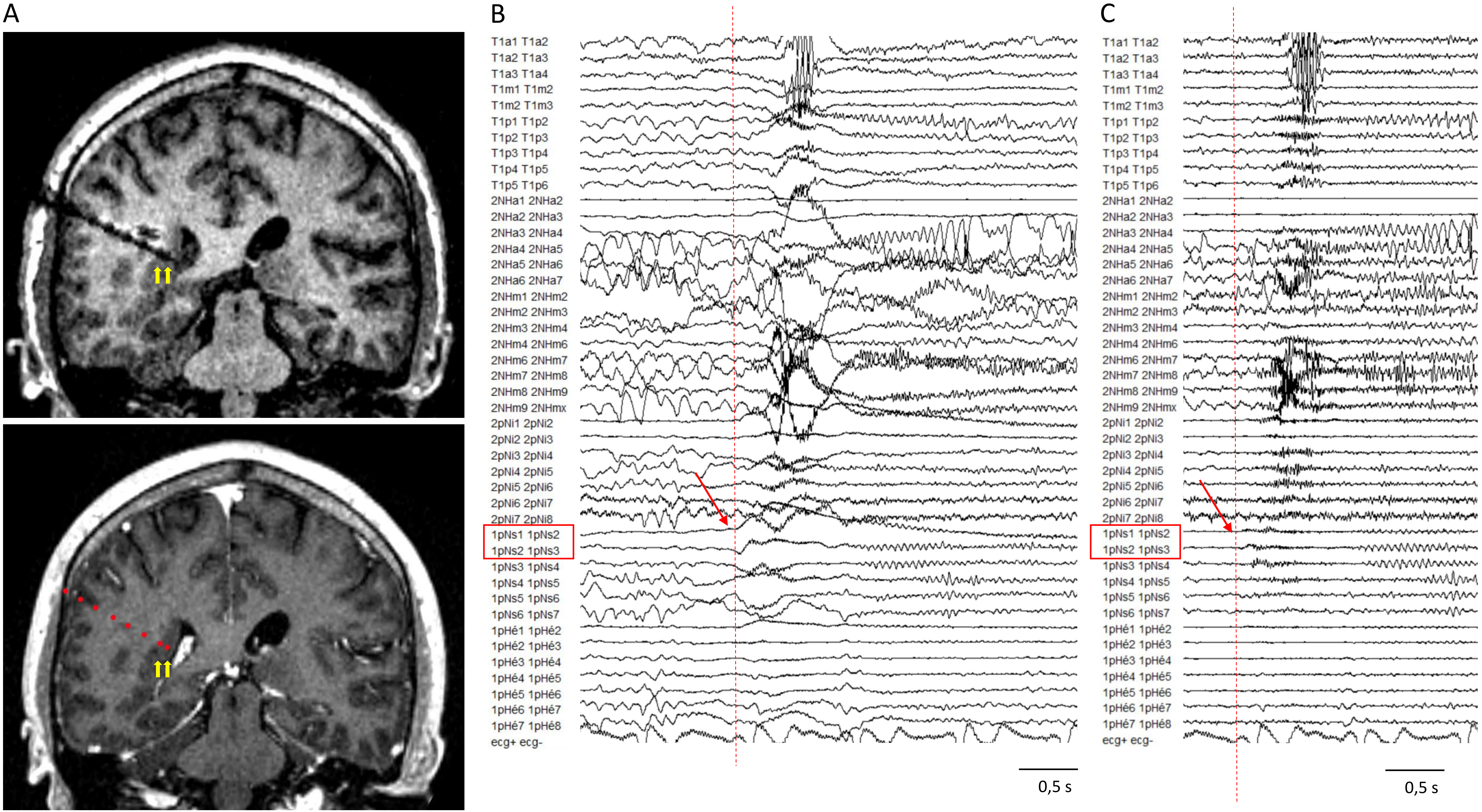

Supplementary Fig. 3B. Patient 2

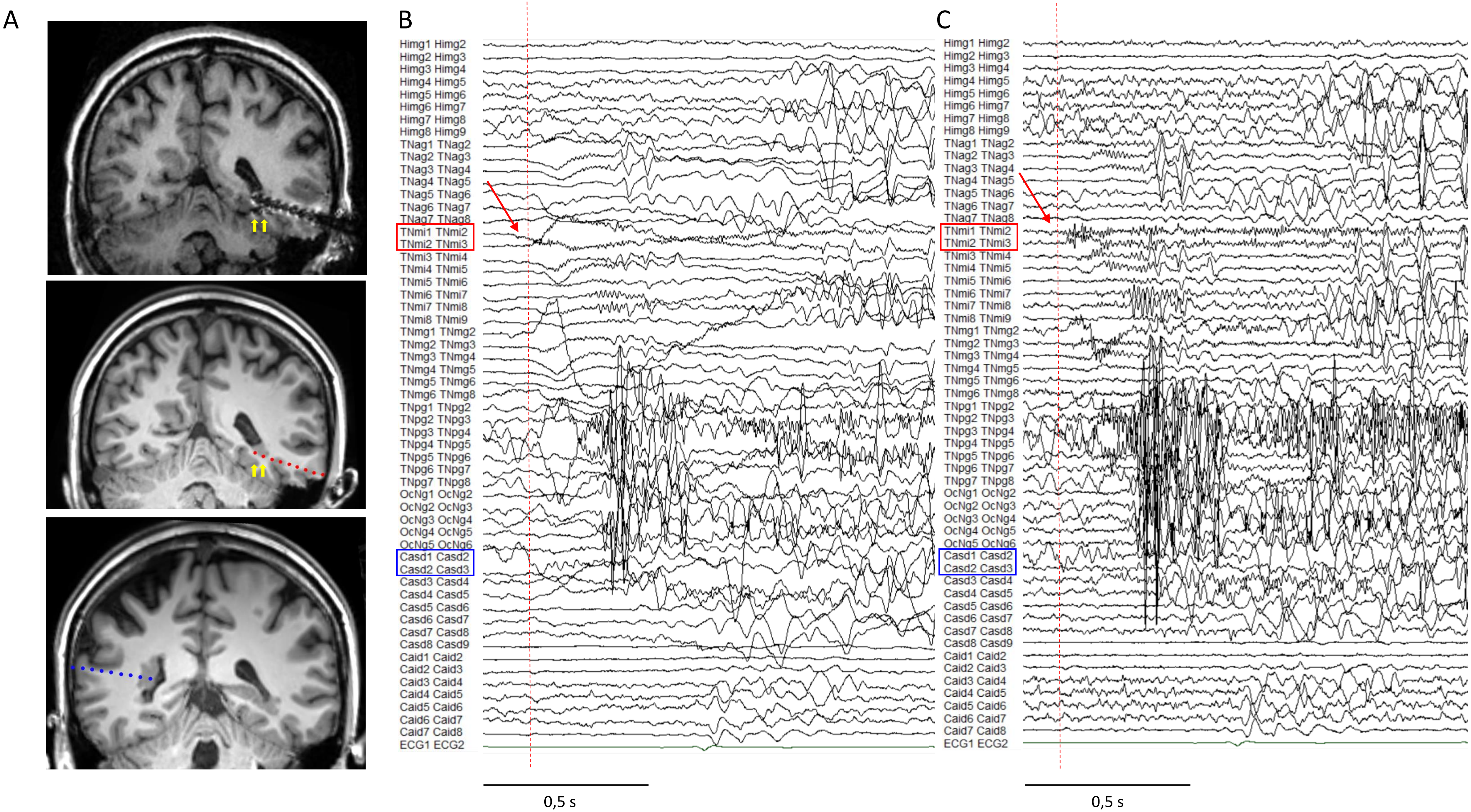

Supplementary Fig. 3C. Patient 3

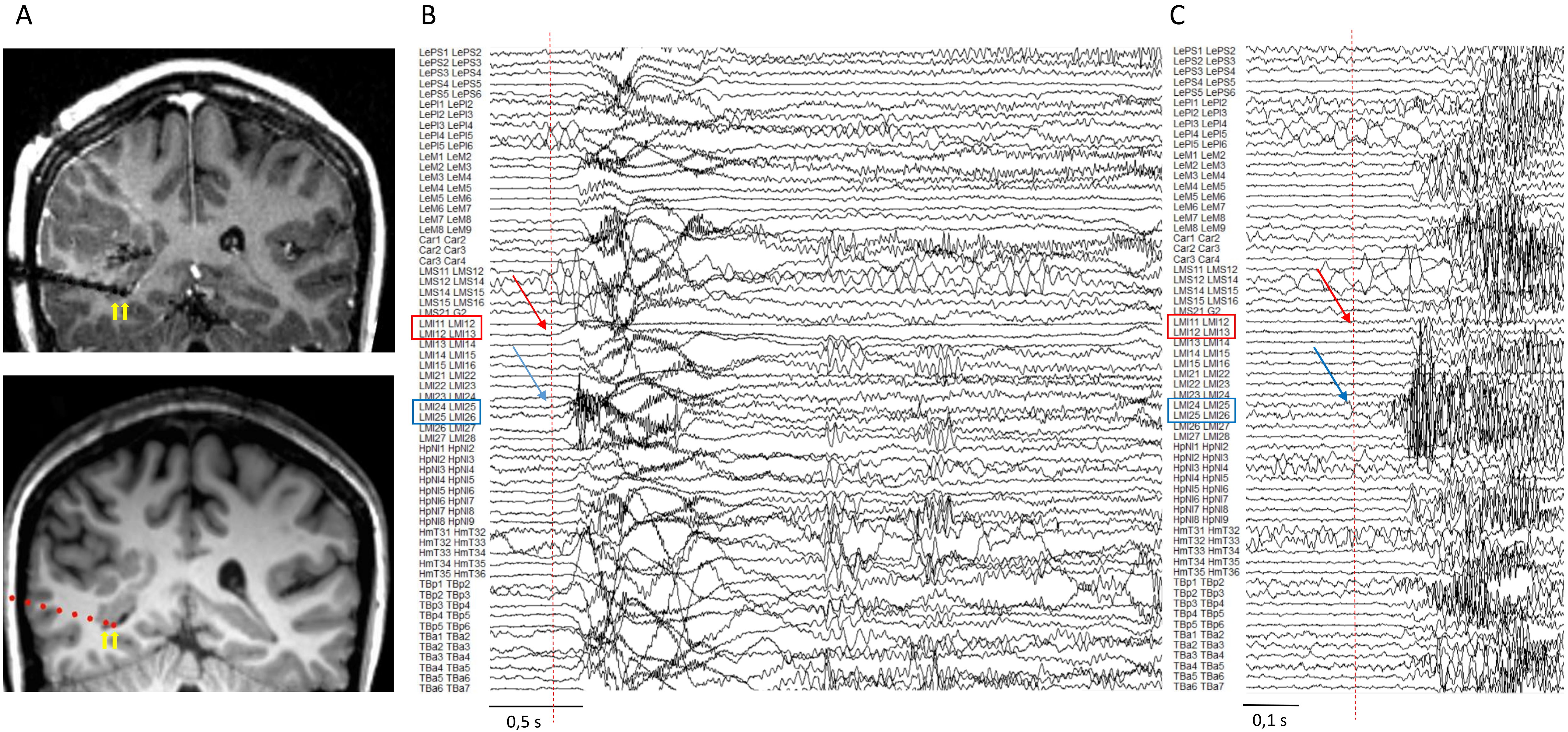

Supplementary Fig. 3D. Patient 3

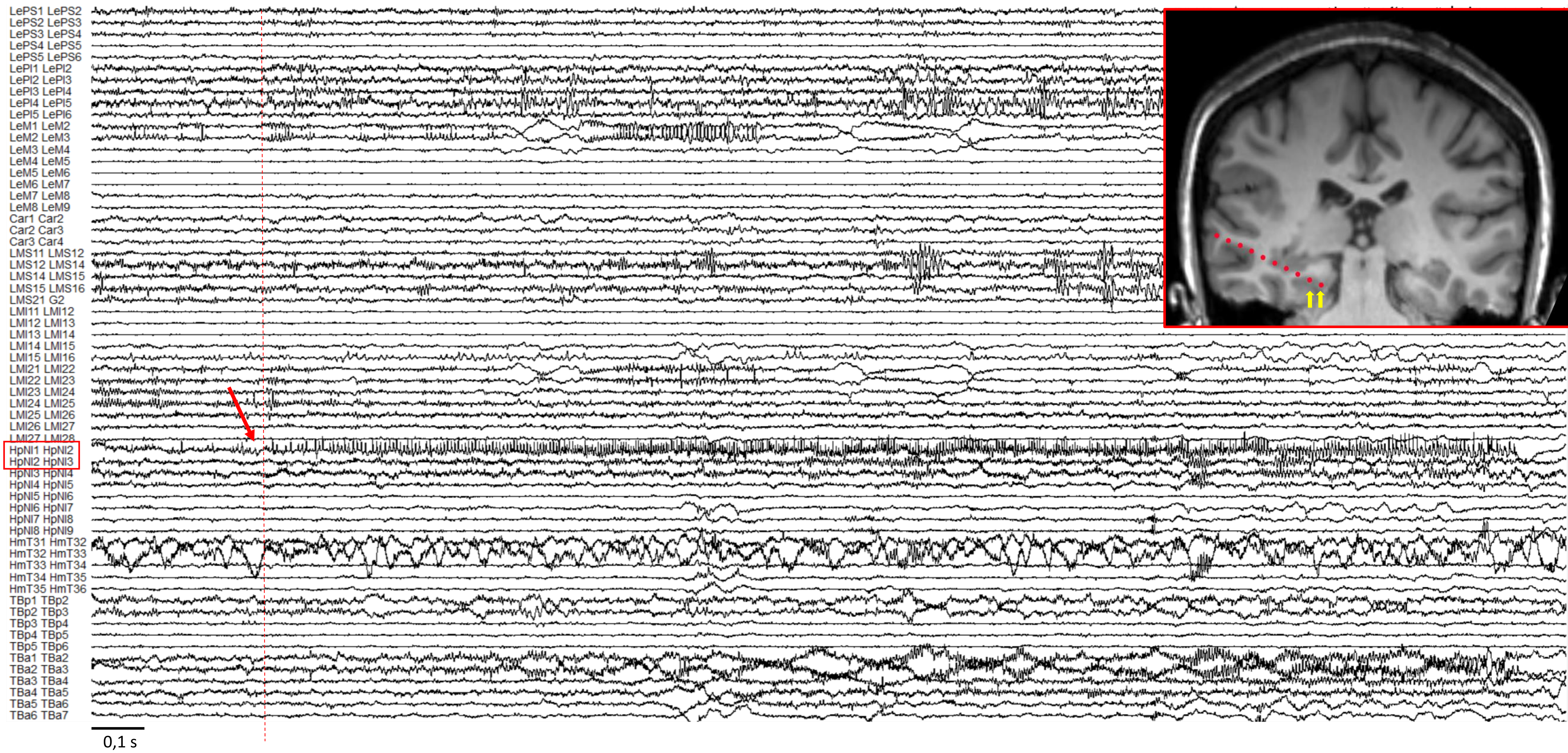

**Supplementary Fig.3: representative SEEG recordings from the three patients**

**Supplementary Fig. 3A. Patient 1**
**A.***Upper panel*: post-implantation MRI showing the trajectory of the electrode exploring the right periventricular nodule. Electrode trajectory is visible despite the magnetic susceptibility artifacts. *Lower panel:* reconstruction of the electrode trajectory superposed to the anatomical preimplantation MRI, allowing a better identification of the anatomical structures targeted by the electrode and avoid anatomical distortions. The yellow arrows indicate the contacts (located in the nodule) primarily involved at seizure start. **B** and **C.** Representative seizure of patient 1 recorded on SEEG. Seizure origin is indicated by the vertical dotted line. Red arrow indicates the contacts from which the electrical origin of seizure is firstly identified. Contacts firstly engaged in seizure start are contoured by a red rectangle. These contacts correspond to those indicate in **A**. The seizure is shown In **B**. high-pass filter: 0,008 Hz; low-pass filter: 1000 Hz; amplitude 300 µV/cm. Bipolar montage. **C.** The same seizure showed after the high-pass filter modification in order to remove slow components and facilitate the identification of the fast component of the ictal discharge. High-pass filter: 20 Hz; low-pass filter: 1000 Hz; amplitude 200 µV/cm. Bipolar montage.

**Supplementary Fig. 3A. Patient 2**
***A****. Upper and middle panels* : post-implantation MRI showing the trajectory of the electrode exploring the left-inferior periventricular nodule of patient 2. *Upper panel:* Electrode trajectory is visible despite the magnetic susceptibility artifacts. Deepest contacts are locate in the inferior part of the periventricular nodule. *Middle panel:* Electrode trajectory is superposed to the preimplantation anatomical MRI, allowing a better identification of targeted anatomical structures avoiding anatomical distortions. The yellow arrows indicate the contacts (located in the nodule) primarily involved at seizure start. *Lower panel:* post-implantation MRI showing the trajectory of the electrode exploring the right periventricular nodule of patient 2 (each blue dot represents an electrode contact). The two deepest contacts are located in the heterotopia. **B** and **C.** Representative recorded seizure of patient 2. Seizure origin is indicated by the vertical dotted line. Red arrow indicates the contacts from which the electrical origin of seizure is firstly identified. Contacts firstly engaged in seizure start are contoured by a red rectangle. These contacts correspond to those indicate in **A** in the upper and middle panel. Note that the electrode exploring the right nodule does not support a primary role of the right nodule in seizure initiation (contacts are contoured by blue rectangle). The seizure is shown In **B**. high-pass filter: 0,160 Hz; low-pass filter: 1000 Hz; Amplitude 250 µV/cm. Bipolar montage. **C.** The same seizure showed after the high-pass and signal amplitude modification in order to remove the slow components. High-pass filter: 20 Hz; low-pass filter: 1000 Hz; Amplitude 110 µV/cm. Bipolar montage.

**Supplementary Fig. 3C. Patient 3**
***A****. Upper panel*: post-implantation MRI showing the trajectory of the electrode exploring the inferior part of the right periventricular nodule of patient 3. *Lower panel:* reconstruction of the electrode trajectory superposed to the anatomical preimplantation MRI, allowing a better identification of targeted anatomical structures avoiding anatomical distortions. The yellow arrows indicate the contacts (located in the nodule) primarily involved at seizure start. **B** and **C.** Representative seizure of patient 3 recorded at SEEG. Seizure start is indicated by the vertical dotted line. The red arrow indicates the contacts from which the electrical origin of seizure is firstly identified. In these patient seizures showed a wider onset with a subtle and precious low voltage activity visible in contacts targeting the inferior part of the right periventricular nodule. These contacts are contoured by a red rectangle. These contacts correspond to those indicate in **A.** A very precocious ictal activity was also visible in another electrode targeting the same nodule and the adjacent polymicrogyric cortex (LMI2, blue rectangles, blue arrows on SEEG trace, anatomical image not shown). The seizure is shown in **B:** High-pass filter: 0,160 Hz; low-pass filter: 1000 Hz; amplitude 250 µV/cm. Bipolar montage. **C.** The same seizure showed before, after the high-pass filter modification in order to remove slow components and facilitate the identification of the fast component of the ictal discharge. High-pass filter: 50 Hz; low-pass filter: 1000 Hz; amplitude 60 µV/cm. Bipolar montage.

**Supplementary Fig. 3D. Patient 3**
***MRI Insert :*** post-implantation MRI showing the trajectory of the electrode exploring the posterior part of the right middle temporal circumvolution and the parahippocampal region. The yellow arrows indicate the contacts (located in the nodule) primarily involved at seizure start.
In patient 3 the posterior right parahippocampal gyrus was the site for generation of focal and short lasting seizures (around the 40 % of the all recorded seizures). Seizure origin is indicated by the vertical dotted line. Red arrow indicates the contacts from which the electrical origin of seizure is firstly identified. Contacts firstly engaged in seizure start are contoured by a red rectangle. high-pass filter: 0,160 Hz; low-pass filter: 1000 Hz; amplitude 400 µV/cm. Bipolar montage.
