## Supplementary Material for "Human periventricular nodular heterotopia shows several interictal epileptic patterns, associated with hyperexcitability of neuronal firing"

### Patient information

**Patient 1**

Patient 1 was investigated for refractory epilepsy starting at the age of 20. Ictal symptoms consisted of vertigo and nausea followed by vocalizations, gestural and alimentary automatisms with loss of consciousness. The patient had at least 20-25 seizures per month and no focal to bilateral tonic-clonic seizures were reported. Brain MRI showed a periventricular nodular heterotopia (PNH) located on the temporal horn of the right ventricle associated with dysplastic overlying cortex. Long-term scalp video-EEG recording detected focal epileptic abnormalities diffusing in the right anterior temporal regions. Seizures started by a fast rhythmic activity in the right temporal region then diffused into bilateral frontal regions with a predominance on the right hemisphere. Surprisingly, the interictal 18FDG-PET scan showed subcortical hyper-metabolism most prominent in the nodular heterotopia. Eight intracerebral electrodes were placed in the right posterior temporal region in order to study the cortical malformation. Implantation was performed at the Department of Neurosurgery of the Pitié-Salpêtrière Hospital using Leksell frame-based stereotaxy. Several spontaneous seizures were recorded during 16 days of continuous recording. They usually originated from the deeper contacts of the two electrodes located into the posterior part of the PNH (Table 1). The most common ictal clinical symptoms were dizziness, staring, loss of consciousness and motor automatisms. Seizures were electrically characterized by the appearance of a slow deflection overlapped with fast rhythms between 115Hz and 160Hz, starting from the posterior part of PNH (Fig. 1J). Electrical stimulation of the posterior part of the nodule promoted the appearance of symptoms described by the patient as typical ictal clinical manifestation (dizziness), giving further evidence that the nodule region was the SOZ.

**Patient 2**

Patient 2 was investigated for focal refractory epilepsy starting at the age of 25. Seizures were described as short tingling sensations of the shoulders and head, hot flushes and vocalizations followed by loss of consciousness. No focal to bilateral seizures were reported. The patient had at least one seizure per day. Surface EEG recordings found abundant interictal epileptic abnormalities in the left posterior temporal or temporo-occipital regions. Seizures started in the left posterior temporal region. Brain MRI revealed bilateral periventricular heterotopia, predominating in the posterior regions, with no abnormalities of adjacent cortical or hippocampal structures. Interictal 18FDG-PET showed that PNHs were isometabolic with the overlying cortex. The ictal SPECT detected an increased blood flow in the left temporal, central and posterior cortical areas, but no modification in the PNH. The patient was then implanted with 8 depth intracranial electrodes investigating the temporo-parietal junctions bilaterally together with the underlying PNHs. Implantation was done using the robotic assistant device ROSA (ROSA® Brain, Medtech, France). Intracerebral interictal abnormalities consisted of epileptic spikes originating from the left posterior nodules then diffusing to the temporal cortex together with large spikes in the left hippocampus. Seizures consisted of tingling sensations of shoulders and trunk, remounting to the head, sometimes followed by forced head rotation to the right, eyelids blinking, small vocalizations and blowing. Electrically, those seizures began with a fast rhythmic activity starting in the anterior part of the nodules, then evolving into large spike-wave discharges initially detectable in the posterior part of the left PNH.

**Patient 3**

In this patient epilepsy started at the age of 14. Epileptic manifestations consisted of vestibular and/or vegetative symptoms often followed by loss of awareness and ipsilateral complex motor automatisms and vocalisations. Seizures could be focal to bilateral. The post-ictal state was characterized by confusion and emotional symptoms (anger and irritability). Seizure duration was of 2-3 minutes. Brain MRI disclosed a PNH located in the posterior part of the right temporal lobe, associated with polymicrogyric abnormalities of the overlying temporal cortex. The long-term EEG video monitoring disclosed a wide focus of IEDs in the right temporal lobe and recorded 3 seizures originating from the right temporal cortex, appearing as a slow deflection followed by a rhythmical activity visible in the right temporal and temporo-frontal leads. Interictal PET found that PNH were isometabolic with normal cortex. Ictal SPECT showed a large hyperperfused area involving the right parieto-temporal cortex, the temporal pole and the PNH. Intracerebral electrodes implantation was performed. The sEEG investigation concluded that the patient suffered from multifocal epilepsy, identifying at least 3 different sites responsible for seizure generation. A first type of focal seizure was characterized by a wide ictal activity start. These seizures usually appeared as diffuse slow deflections, associated with rapid low voltage fast activity, visible on multiple electrodes. Despite this wide appearance, some contacts were activated slightly earlier, especially those located in the subependymal nodules. About these nodules, the most inferior and posterior part of the right nodular lesion was the one showing the earliest component of the ictal discharge. The adjacent cortices (apparently normal on MRI), the posterior temporo-basal cortex, the temporo-parietal junction and the lateral right temporal cortex can sometimes participate. A second type of focal seizure, started from the parahippocampal region. These seizures could remain very focal for a long time (or even exclusively), or propagate in the temporo-basal cortex, before spreading to other regions. The hippocampus is never an initiator and is usually involved in the later stages of seizure development. Seizures starting from the PNH were often either asymptomatic or mildly symptomatic. When symptomatic, seizures were longer in duration and often they involved the temporo-basal cortex or the hippocampus. A less common type of seizures was characterized by a progressive intensification of interictal discharges, which subsequently formed into a seizure, but without being clinically symptomatic. This latter type of seizures was more frequent during sleep.
